# Multicellular immune hubs and their organization in MMRd and MMRp colorectal cancer

**DOI:** 10.1101/2021.01.30.426796

**Authors:** Karin Pelka, Matan Hofree, Jonathan Chen, Siranush Sarkizova, Joshua D. Pirl, Vjola Jorgji, Alborz Bejnood, Danielle Dionne, William H. Ge, Katherine H. Xu, Sherry X. Chao, Daniel R. Zollinger, David J. Lieb, Jason W. Reeves, Christopher A. Fuhrman, Margaret L. Hoang, Toni Delorey, Lan T. Nguyen, Julia Waldman, Max Klapholz, Isaac Wakiro, Ofir Cohen, Christopher S. Smillie, Michael S. Cuoco, Jingyi Wu, Mei-ju Su, Jason Yeung, Brinda Vijaykumar, Angela M. Magnuson, Natasha Asinovski, Tabea Moll, Max N. Goder-Reiser, Anise S. Applebaum, Lauren K. Brais, Laura K. DelloStritto, Sarah L. Denning, Susannah T. Phillips, Emma K. Hill, Julia K. Meehan, Dennie T. Frederick, Tatyana Sharova, Abhay Kanodia, Ellen Z. Todres, Judit Jané-Valbuena, Moshe Biton, Benjamin Izar, Conner D. Lambden, Thomas E. Clancy, Ronald Bleday, Nelya Melnitchouk, Jennifer Irani, Hiroko Kunitake, David L. Berger, Amitabh Srivastava, Jason L. Hornick, Shuji Ogino, Asaf Rotem, Sébastien Vigneau, Bruce E. Johnson, Ryan Corcoran, Arlene H. Sharpe, Vijay K. Kuchroo, Kimmie Ng, Marios Giannakis, Linda T. Nieman, Genevieve M. Boland, Andrew J. Aguirre, Ana C. Anderson, Orit Rozenblatt-Rosen, Aviv Regev, Nir Hacohen

**Affiliations:** Broad Institute of Massachusetts Institute of Technology (MIT) and Harvard, Cambridge, MA, USA; Massachusetts General Hospital Cancer Center, Harvard Medical School (HMS), Boston, MA, USA; Klarman Cell Observatory, Broad Institute of MIT and Harvard, Cambridge, MA, USA; Department of Pathology, Massachusetts General Hospital, Boston, MA, USA; Department of Biomedical Informatics, Harvard Medical School, Boston, MA, USA; NanoString Technologies Inc., Seattle, WA, USA; Evergrande Center for Immunologic Diseases, Harvard Medical School and Brigham and Women’s Hospital, Boston, MA, USA; Center for Cancer Genomics, Department of Medical Oncology, Dana-Farber Cancer Institute, Boston, MA, USA; Department of Medical Oncology, Dana-Farber Cancer Institute, Boston, MA, USA; Department of Immunology, Harvard Medical School, Boston, MA, USA; Clinical Research Center, Massachusetts General Hospital, Boston, MA, USA; Clinical Research Center, Dana-Farber Cancer Institute, Boston, MA, USA; Department of Molecular Biology, Massachusetts General Hospital, Boston, MA, USA; Columbia Center for Translational Immunology, New York, NY; Columbia University Medical Center, Division of Hematology and Oncology, New York, NY; Program for Mathematical Genomics, Columbia University, New York, NY; Department of Surgery, Brigham and Women’s Hospital, Boston, MA, USA; Department of Surgery, Massachusetts General Hospital, Boston, MA, USA; Department of Pathology, Brigham and Women’s Hospital, Boston, MA, USA; Department of Immunology, Blavatnik Institute, Harvard Medical School, Boston, MA, USA; Department of Medicine, Harvard Medical School, Boston, MA, USA; Howard Hughes Medical Institute and Koch Institute for Integrative Cancer Research, Department of Biology, Massachusetts Institute of Technology, Cambridge, MA, USA

**Author notes:** Genentech, 1 DNA Way, South San Francisco, CA. These authors contributed equally to this work (Karin Pelka, Matan Hofree, Jonathan Chen). These authors contributed equally to this work (Aviv Regev, Nir Hacohen). Corresponding author (A.C.A.); (O.R.); (A.R.); (N.H.).

**Keywords:** Colorectal cancer, anti-tumor immunity, mismatch repair-deficient, mismatch repair-proficient, MSS, MSI, spatial, scRNA-seq, cell-cell interactions

## Abstract

Immune responses to cancer are highly variable, with mismatch repair-deficient (MMRd) tumors exhibiting more anti-tumor immunity than mismatch repair-proficient (MMRp) tumors. To understand the rules governing these varied responses, we transcriptionally profiled 371,223 cells from colorectal tumors and adjacent normal tissues of 28 MMRp and 34 MMRd patients. Analysis of 88 cell subsets and their 204 associated gene expression programs revealed extensive transcriptional and spatial remodeling across tumors. To discover hubs of interacting malignant and immune cells, we identified expression programs in different cell types that co-varied across patient tumors and used spatial profiling to localize coordinated programs. We discovered a myeloid cell-attracting hub at the tumor-luminal interface associated with tissue damage, and an MMRd-enriched immune hub within the tumor, with activated T cells together with malignant and myeloid cells expressing T-cell-attracting chemokines. By identifying interacting cellular programs, we thus reveal the logic underlying spatially organized immune-malignant cell networks.

## Introduction

Almost all tumors are infiltrated with immune cells, but the types of immune responses and their effects on tumor growth, metastasis and death, vary greatly between different cancers and individual tumors^1^. Immune responses arise from the interactions between immune, malignant, stromal and other cell types in the tumor microenvironment^2, 3^, as well as due to contact with microbes and their ligands^3, 4^. Which of the numerous cell subsets in a tumor contribute to the response, how their interactions are regulated, and how they are spatially organized within tumors remains poorly understood^5, 6^.

Colorectal tumors show a large dynamic range of immune responsiveness, with a striking difference between two genetically distinct subtypes^7, 8^. Mismatch repair-deficient (MMRd) colorectal tumors have a high mutational burden, often contain cytotoxic T cell infiltrates, and have a ∼50% response rate to immune checkpoint blockade, while mismatch repair-proficient (MMRp) tumors have a low mutational burden and are largely unresponsive to immunotherapy^9–12^. MMR deficiency is most commonly caused by promoter methylation of MLH1, which is associated with BRAF mutations and tends to occur in older patients^7^. Germline mutations (such as in Lynch syndrome) or somatic alterations can also cause MMR deficiency and are associated with TP53 or KRAS mutations^13, 14^.

Transcriptional profiles of bulk tumors^15, 16^ or single cells^17–20^, have been used to classify colorectal tumors into subtypes, define their cellular composition, and infer interaction networks between cell types. However, these studies focused on discrete cell clusters, and thus did not capture the full spectrum of transcriptional programs, which can exist as continuous gradients of program activities within or across clusters^21, 22^. Furthermore, such studies predicted interactions between cell type clusters based on the expression of receptor-ligand pairs, but not by joint functional changes across cell types. Recently, imaging-based studies have highlighted cellular interaction networks based on the recurrent co-localization of different cells in neighborhoods^23^, but were limited by the number of pre-selected markers that resolve key cell types but not their finer features or comprehensive molecular profiles.

Here, we developed a systematic approach to discover cell types, their underlying programs, and cellular communities based on single cell RNA-seq (scRNA-seq) profiles and applied it to study the distinguishing features of human MMRd and MMRp colorectal cancer. We focused on understanding the immune responses within these tumors, and did not consider tumor genetics or neoantigens in this study. We identified 88 cell subsets across immune, stromal and malignant cells, and 204 associated gene expression programs. We revealed multicellular interaction networks based on co-variation in the activity of expression programs in different cell subsets across patients, and imaged key molecules for predicted cell subsets and programs to localize these interaction networks in matched tissue from the same patient cohort. We found stromal remodeling that resulted in the reduction of BMP-producing fibroblasts in MMRd tumors and the mis-localization of fibroblast-derived stem cell niche factors throughout the tumor. We discovered an inflammatory interaction network of malignant cells, monocytes, fibroblasts, and neutrophils at the luminal margin of primary MMRd and MMRp tumors, and MMRd-specific hotspots of immune activity comprised of chemokine-expressing malignant and non-malignant cells adjacent to activated T cells. Our study demonstrates a path to discovering multicellular interaction networks that underlie immunologic and tumorigenic processes in human cancer.

## Results

### A comprehensive atlas of cell subsets, programs, and multicellular interaction networks in MMRd and MMRp colorectal cancer

To discover how malignant, immune, and stromal cells interact in MMRd and MMRp CRC, we collected and analyzed primary untreated tumors from 34 MMRd and 28 MMRp patients (with an additional lesion collected for 2 patients) as well as adjacent normal colon tissue for most patients (Figure 1A, **Supplemental Table 1A**). We performed droplet-based scRNA-seq on dissociated fresh tissues, retaining 371,223 high quality cells (**Methods**), including 168,672 epithelial (non-malignant and malignant), 187,094 immune, and 15,457 stromal cells (Supplemental Figure 1A,B).

**Figure 1.**
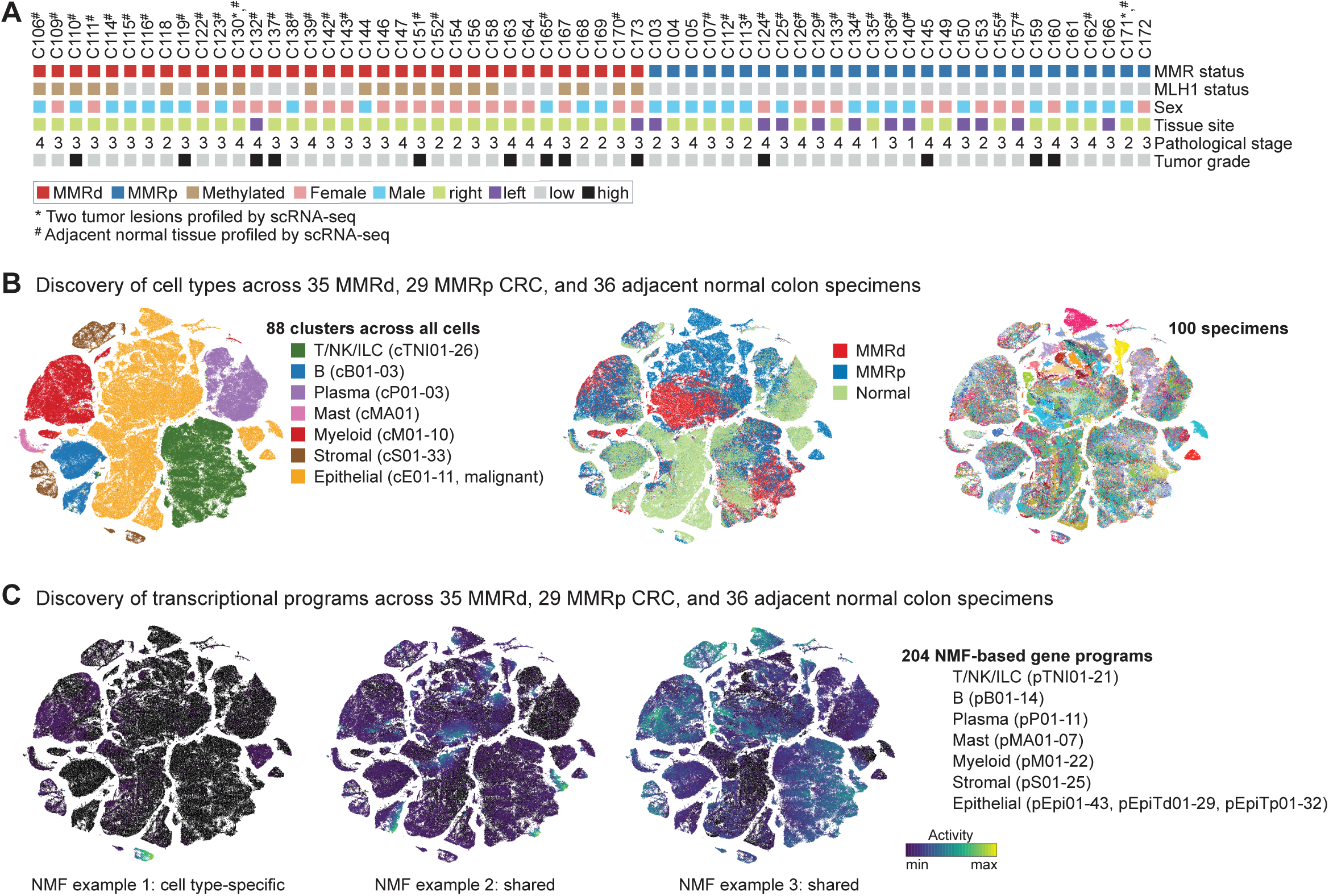
Patient cohort and atlas of cell subsets and programs in MMRd and MMRp colorectal cancer. (**A**) MMR status (34 MMRd and 28 MMRp), and key clinical characteristics of primary untreated CRC specimens. (**B**) tSNE visualization of 371,223 cells based on scRNA-seq expression profiles and colored by major cell partitions (left), by tissue type (middle), or patient specimen (right). (**C**) In addition to clusters, 204 NMF-based gene programs were derived across all cell partitions, among them both cell type-specific and shared transcriptional programs, illustrated by programs pS02 (Fibro matrix/stem cell niche), pTNI03 (proliferation), pEpi30 (ISGs). See also Supplemental Figure 1 and **Supplemental Table 1**.

We defined cell subsets and transcriptional programs by a two-step graph-clustering approach: first, we clustered all cells into 7 major partitions (T/NK/ILC, B, plasma, mast, myeloid, stroma/endothelial and epithelial); second, within each partition, we derived clusters (prefix ‘c’) and transcriptional programs (sets of genes with co-varying expression, prefix ‘p’) using consensus non-negative matrix factorization (NMF)^22, 24^ (Figure 1B,C; **Supplemental Table 2-4; Methods**). *De novo* identification of programs by NMF enabled several key analyses: (1) simultaneous identification of programs shared across multiple cell types (e.g. proliferation, metabolic and immune programs), specific to a cell type (e.g. pDC program), and/or expressed in continuous gradients within or across cell clusters; (2) finding of shared biological properties of malignant cells across patients despite strong patient-specific transcriptional states^25, 26^; and (3) identification of co-varying programs across multiple tumor specimens to find networks of coordinated cells or states that reflect cell interactions or response to a common trigger.

### Remodeling of the immune cell compartment in MMRd and MMRp colorectal tumors

To understand the basis for differential immune responses in CRC, we first determined and compared the immune cell composition of MMRd and MMRp CRC and normal colon tissue, finding dramatic remodeling between tumor and normal tissue and between MMRd and MMRp tumors. Specifically, 37 of 43 immune cell clusters (manually curated cluster markers in Supplemental Figure 2A) were differentially abundant as a fraction of all immune cells between tumor (either MMRd or MMRp) and normal colon tissue (Figure 2A, Supplemental Figure 2B, **Supplemental Table 2**). Tumors were depleted of IgA-producing plasma cells, B cells, *IL7R*+ T cells and *γδ*-like T cells, and enriched with Tregs, monocytes, macrophages and likely neutrophils relative to adjacent normal colon (Figure 2A).

**Figure 2.**
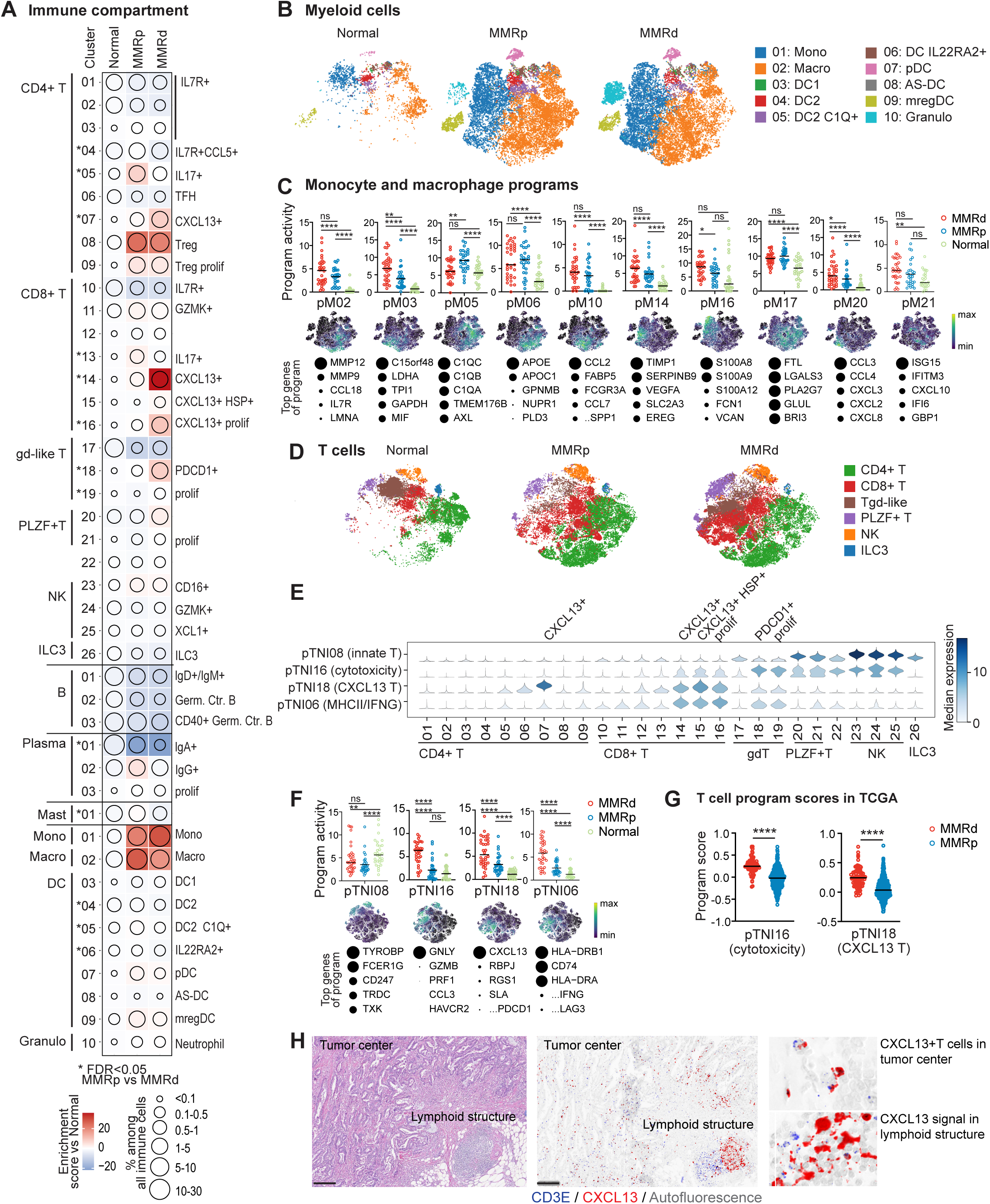
The immune compartment in MMRd and MMRp CRC. (**A**) Changes in immune cell clusters in MMRp and MMRd tumors relative to adjacent normal tissue, showing frequency of immune cells (dot size) or enrichment/depletion (Pearson residual, colored squares). Clusters with differences in frequency between MMRp and MMRd tumors with Kruskal-Wallis FDR<0.05 are marked with *. (**B**) tSNE plot of myeloid cells in all normal and tumor samples. (**C**) Activities of selected myeloid transcriptional programs with high activities in monocytes and macrophages, shown for both tumor types and normal tissue. Each dot indicates the 75th percentile of the program activity in the myeloid cells of one patient specimen. GLME (generalized linear mixed model) FDR is reported as: **** for ≤0.0001, *** for ≤0.001, **≤0.01, *≤0.05, ns for >0.05. tSNEs below show program activities within the myeloid compartment. For each program, the top genes are listed below, with circle size indicating the relative weight of each gene within the program (reflecting uniqueness relative to other programs; scaled to [0,1] range). (**D**) tSNE visualization of the T/NK/ILC partition colored by major cell subsets. (**E**) Activities of programs pTNI08, pTNI16, pTNI18, and pTNI06 within each of the T/NK/ILC clusters. (**F**) Activities of pTNI08, pTNI16, pTNI18, and pTNI06 displayed as in (C), with tSNE of the T/NK/ILC compartment. GLME (generalized linear mixed model) FDR is reported as: **** for ≤0.0001, *** for ≤0.001, **≤0.01, *≤0.05, ns for >0.05 (**G**) Score for gene signatures of pTNI16 and pTNI18 in bulk RNA-seq from TCGA-CRC (COADREAD) patient specimens. Mann–Whitney–Wilcoxon test **** for p≤0.0001. (**H**) Localization of *CXCL13*+ T cells. Left, H&E showing tumor center and lymphoid structure. Middle, same area stained for *CD3E* and *CXCL13* by multiplex RNA ISH. Right, *CXCL13*+ *CD3E*+ T cells are found in the tumor center (top) and *CXCL13*+ cells are *CD3E-* in lymphoid structures (bottom). Scale bar: 200um. See also Supplemental Figure 2 and **Supplemental Table 2**.

In particular, there was a significant expansion of monocytes and macrophages in tumors compared to normal samples (Figure 2A,B). Monocytes and macrophages upregulated several tumor-specific NMF-derived transcriptional programs (Figure 2B,C), that were characterized by genes that can amplify inflammation (*MMP12* and *MMP9* in pM02), recruit myeloid cells (chemokines *CCL2* and *CCL7* in pM10), stimulate growth (growth factors *VEGFA* and *EREG* in pM14), and resolve inflammation (*APOE* in pM06). MMRd cells showed higher activities of programs with genes in glycolysis (pM03), immune-activating alarmins such as *S100A8/9/12* (pM16) and chemokines that attract monocytes and neutrophils (pM20). Overall, monocytes and macrophages were remodeled in tumors, and expressed more immune-activating programs in MMRd tumors.

### T cell compartment differences between MMRd and MMRp tumors

The predominant change in the immune composition of MMRd versus MMRp tumors was in the T cell compartment (Figure 2A,D). Among the clusters enriched in MMRd tumors were *CXCL13*+ T cells and *PDCD1*+ *γδ*-like T cells, while IL17+ T cells were enriched in MMRp tumors (Figure 2A, marked with * next to cluster number, Supplemental Figure 2B). *CXCL13* in T cells has been noted in other melanoma and CRC single cell studies^17, 18, 27^, and has recently emerged as a marker of human tumor-reactive CD8+ T cells and response to immunotherapy^28–30^. Thus, we hypothesize that anti-tumor T cell immunity may have developed in many MMRd tumors, but very rarely in MMRp tumors (Supplemental Figure 2B).

Programs enriched in MMRd versus MMRp T cells (**Supplemental Table 2E**) included two programs (pTNI18 with *CXCL13, PDCD1, TOX*; pTNI06 with MHC Class II gene, *IFNG* and *LAG3*) with high and moderate activity in TCR*ɑβ* and TCR*γδ*-like T cells respectively, and one cytotoxicity program (pTNI16) shared among CD8+, *γδ*-like, *PLZF*+ T cells and NK cells. *PLZF*+ T cells and NK/ILC3 cells were also selectively marked by expression of an innate/T program (pTNI08) that was reduced in both MMRd and MMRp tumors compared to normal tissue (Figure 2E,F). Using bulk RNA-seq datasets, we confirmed the higher MMRd activity of the *CXCL13* and cytotoxicity programs (which can be attributed only to the T/NK/ILC partition, allowing us to analyze bulk data Supplemental Figure 2C) in three external CRC cohorts (Figure 2G, Supplemental Figure 2D;^13, 31, 32^). Thus, in MMRd tumors, subsets of T and NK cells acquire cytolytic properties (*GNLY*, *GZMB*, *PRF1*), and T cells acquire exhaustion markers associated with (e.g., *PDCD1, TOX, LAG3, HAVCR2*).

### CXCL13+ T cells localize within MMRd tumors

Given our observed enrichment of *CXCL13*+ T cells in MMRd tumors, and their previous association with immunotherapy responses as well as localization to tertiary lymphoid structures (TLS) in lung^28^, we stained tissue sections from our cohort with RNA probes targeting *CXCL13* and *CD3E*. We found abundant *CXCL13*+ T cells throughout the tumor of MMRd specimens, outside of TLS (Figure 2H), which are usually found at the invasive border^33^, whereas TLS-associated CXCL13 expression was largely in non-T (CD3E-negative) cells in a reticular pattern, consistent with reports of stromal and follicular dendritic cells as sources of *CXCL13* in TLS^34^. In summary, *CXCL13*-expressing conventional CD4+ and CD8+ T cells were localized outside of lymphoid structures, but in very close proximity to carcinoma cells, consistent with effector activity.

### Highly altered endothelial cells in both MMRd and MMRp tumors

The stromal compartment was remodeled in both tumor types (Figure 3A,B; Supplemental Figure 3A-C; **Supplemental Table 3**), with an increase in endothelial cells and pericytes as a fraction of stromal cells (Figure 3C) and a reduction in lymphatic endothelial cells as a fraction of endothelial cells in tumor versus normal (Figure 3B). Along with one cluster shared between tumor and normal samples, we found 8 tumor-specific clusters of endothelial cells, with no significant differences between MMRd and MMRp tumors. Quantifying the similarity between endothelial clusters in tumor versus normal colon (using partition-based graph abstraction, PAGA^35^), we found altered versions of arterial and venous cells and several clusters that did not map back to normal cells, such as tip cells and proliferating cells (Figure 3D). Interestingly, these proliferating endothelial cells expressed *HIF1A* and *CSF3* (Supplemental Figure 3A), suggesting metabolic and inflammatory changes.

**Figure 3.**
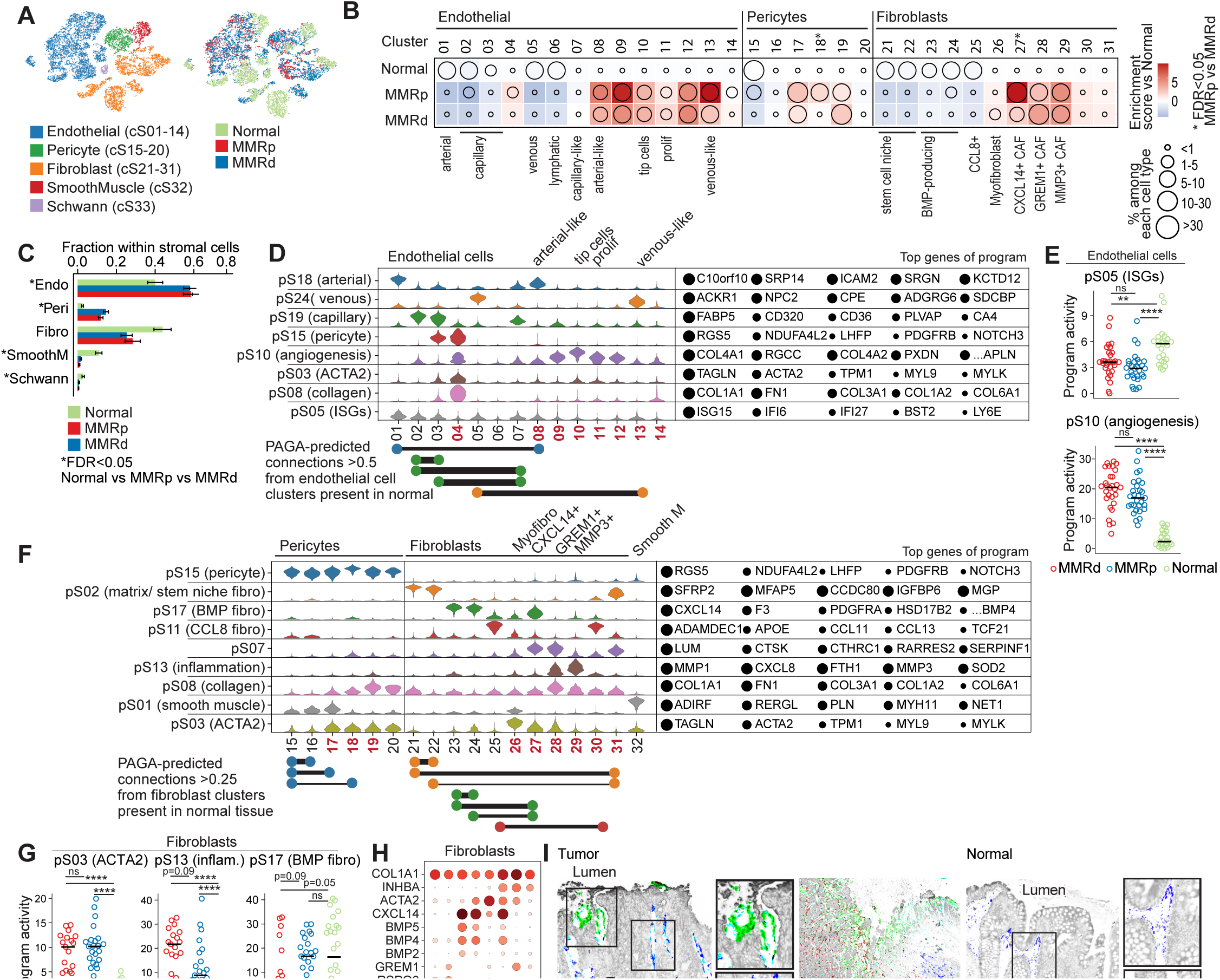
Stromal remodeling in MMRd and MMRp CRC. (**A**) tSNE plot of stromal cells in all normal and tumor samples. (**B**) Changes in endothelial, pericyte, and fibroblast subsets within their respective compartments in MMRp and MMRd tumors relative to adjacent normal tissue, showing frequency (dot size) or enrichment/depletion (Pearson residual, colored squares). Clusters with differences in frequency between MMRp and MMRd tumors with Kruskal-Wallis FDR<0.05 are marked with *. Note that cS30 and cS31 were found predominantly in two patients in which the tumor grew underneath non-neoplastic tissue. (**C**) Fraction of stromal cell subsets per tissue type. Clusters with differences in frequency between normal and tumor with Kruskal-Wallis FDR<0.05 are marked with *. (**D**) Activities of selected programs in each of the endothelial cell clusters. Tumor-enriched clusters are indicated in bold red. Top program genes are listed to the right, with circle size indicating the weight of each gene in the program. Key edges (connectivity) between normal clusters or with tumor-associated clusters (weights >0.5, identified by PAGA) are shown below and colors are matched to programs with high activity in the respective clusters. (**E**) Activity of pS05 (ISG) and pS10 (angiogenesis) in all tumor and normal samples. Each point indicates 75th percentile of the program activity per patient specimen in the endothelial cells. GLME (generalized linear mixed model) FDR is reported as: **** for ≤0.0001, *** for ≤0.001, **≤0.01, *≤0.05, ns for >0.05. (**F**) Activities of selected programs in fibroblast and pericyte subtypes. Tumor-enriched clusters are indicated in bold red. Top program genes are listed to the right, with circle size indicating the weight of each gene in the program. Key edges (connectivity) between normal or with tumor-associated clusters with weights >0.25 (as identified by PAGA) are shown below. Colors are matched to programs with high activity in the respective clusters. (**G**) Activities of pS03 (ACTA2), pS13 (inflammation), and pS17 (BMP fibro) in fibroblasts and pS03, pS13 in pericytes, shown as in (E). (**H**) Dot plot showing geometric mean expression (log(TP10K+1)) and frequency (dot size) of key genes in selected fibroblast subtypes. INHBA distinguishes CAFs from fibroblasts in normal tissue. Tumor-enriched clusters are indicated in bold red. (**I**) Representative multiplex RNA ISH/IF images of tumor show MMP3+ fibroblasts at the luminal surface around dilated vessels (left image). Like CXCL14+ cells (left and middle), GREM1+ cells line malignant cells (middle), but are additionally found in stromal bands reaching far into the tumor center (left). In normal (right) only *CXCL14+* cells line epithelial cells; *GREM1* signal is restricted to in and below the muscularis mucosa (right). Scale bar: 100um (except where specifically annotated). (**J**) Quantification of *CXCL14*+, *GREM1*+ and *MMP3*+ CAFs among all *COL1A1*/*COL1A2*+ fibroblasts based on whole slide scans of 5 MMRd and 4 MMRp CRC specimens from panel (I), Mann-Whitney-Wilcoxon test. Rightmost graph, *MMP3*+ cells among all *COL1A1*/*COL1A2*+ cells outside or inside of the luminal margin (defined as ≤ 360 um from the luminal border of the tumor), Wilcoxon matched-pairs signed rank test. Note 8 samples included at right because clinical paraffin block preparation required removal of luminal margin from one sample. (**K**) Score for gene signatures of *CXCL14*+ CAFs, *GREM1*+ CAFs, *MMP3*+ CAFs, and all fibroblasts in bulk RNA-seq from TCGA-CRC (COADREAD) patient specimens. Mann–Whitney–Wilcoxon test **** for p≤0.0001, *** for ≤0.001, **≤0.01, *≤0.05, ns for >0.05. (**L**) Multiplex RNA ISH/IF shows *RSPO3* signal is restricted to the crypt base in normal colon tissue (right image) but ascends far into the tumor (left). The stainings were performed on consecutive sections to the stainings shown in (I). See also Supplemental Figure 3 and **Supplemental Table 3**.

Program pS10, which contains basement membrane collagens, pro-angiogenic molecules and a tip cell marker (**Supplemental Table 3**), was upregulated across all tumor-specific clusters, whereas a program of interferon stimulated genes (ISGs)/antigen presentation (pS05) was repressed (Figure 3D,E), as observed previously^18^. Thus, endothelial cells are highly altered in tumors, with more angiogenesis program activity and changes in immune-related gene expression.

### Inflammatory fibroblasts localize to the luminal surface of tumors

Fibroblasts partitioned into 11 subsets, with 6 predominant in tumor and 5 in normal colon samples (Figure 3B). Analogous to the previously described myCAFs^36–38,^ 3 cancer-associated fibroblast subsets (cS26-28) (and tumor pericytes) expressed a contractile program (pS03) that included smooth muscle actin (*ACTA2*) (Figure 3F,G, **Supplemental Table 3**), with one subset (cS26; myofibroblasts) expressing it very highly along with the smooth muscle program (pS01) which was shared with smooth muscle cells and pericytes (Figure 3F).

Two CAF subsets (cS28,29) expressed an inflammatory program (pS13) (Figure 3F, **Supplemental Table 3**) in both tumor types, with higher activity in MMRd tumors (Figure 3G and **Supplemental Table 3**). This program, mirroring those of previously described inflammatory CAFs (iCAFs)^36–38^ and inflammatory fibroblasts in ulcerative colitis^39–41^, included tissue remodeling factors (*MMP2*, *MMP3*) and neutrophil-attracting chemokines (*CXCL8*, *CXCL1*). Tissue staining for *MMP3* and the ubiquitous fibroblast marker *COL1A1* (Figure 3H) in 8 CRC specimens (4 MMRd, 4 MMRp), revealed that these highly inflammatory fibroblasts were strongly enriched around dilated blood vessels at the colonic luminal margin (LM) of both MMRd and MMRp tumors (Figure 3I,J, Supplemental Figure 3D).

### *BMP*-expressing CAFs are reduced in MMRd CRC, whereas CAF-derived stem cell niche factors are abnormally present throughout tumors

To further understand the functional alterations in CAFs and their potential impact, we compared the CAFs to fibroblasts from adjacent normal colon tissue based on shared programs and PAGA-based similarity between clusters^35^ (Figure 3F). We identified a CAF equivalent (cS27) of BMP-expressing fibroblasts, cells that line normal colon epithelial cells and drive the differentiation of epithelial cells through WNT inhibition via BMPs and WNT antagonists such as *FRZB*. These may correspond to PDGFRA-high subset of telocytes in the small intestine^42^. The BMP-expressing CAFs were distinguished from other CAF subsets by *CXCL14* expression (Figure 3H), and *CXCL14*+ fibroblasts lined the epithelium in both normal and tumor specimens (Figure 3I). A previous bulk RNA-seq study reported reduced *CXCL14* expression in MMRd vs. MMRp CRC, but suggested this was due to differential expression in malignant epithelial cells^16^. While there is a significant, but modest (1.25-fold reduction) change in *CXCL14* expression of MMRd vs. MMRp malignant cells, they rarely expressed *CXCL14* (∼9.2% of MMRp and ∼1.5% of MMRd malignant cells), with one exception (MMRp patient C103, Supplemental Figure 3E). Instead, MMRd patients (as well as MMRp patient C107 who had high T cell activity) had reduced *CXCL14*+ CAFs (Figure 3B), which we confirmed in imaging-based quantification (Figure 3J) and external cohorts^31, 32^ (Figure 3K; Supplemental Figure 3F).

CAFs also contributed expression of stem cell niche factors, such as RSPO3 and GREM1, which were broadly expressed throughout tumors (Figure 3I **left**, **3L left**), in contrast to their crypt-associated expression in normal tissue (Figure 3I **right, 3L right**). Specifically, in non-neoplastic tissue, RSPO3 and GREM1 expression is strictly limited to areas below the bottom of the crypt^43–45^ most prominently along a distribution similar to that of the muscularis mucosa (Supplemental Figure 3G), as described previously^44, 46, 47^. In contrast, *GREM1*+ and *RSPO3*+ cells (Figure 3I,L) were found in stromal bands that reached far upward from the base into the tumor body. In MMRd specimens, these cells also occupied positions similar to the epithelial cell-lining *CXCL14*+ *BMP*-expressing fibroblasts (Figure 3I**, middle image**). High expression of *RSPO3* drives tumor growth and can arise from *PTPRK-RSPO3* fusion events in a small fraction of human CRC^48, 49^. Our data suggest that perhaps a more common mechanism to increase access to stem cell niche factors, like *RSPO3,* occurs via spatial redistribution of stromal cells, especially CAFs.

### Malignant cells are actively engaged in the immune response

Since malignant cells typically group by patient (in contrast to normal epithelial cells that cluster by cell subset) (Figure 4A), it can be more challenging to identify their shared properties. We therefore derived (**Methods**) and analyzed the activities of 43 expression programs in malignant cells (denoted pEpi*; Figure 4B; Supplemental Figure 4A,B; **Supplemental Table 4**), which were not specific to single patients. We also categorized malignant cells based on similarity to normal colon epithelial cell subtypes to better understand their functional properties (Figure 4C, Supplemental Figure 4C,D, **Methods**).

**Figure 4:**
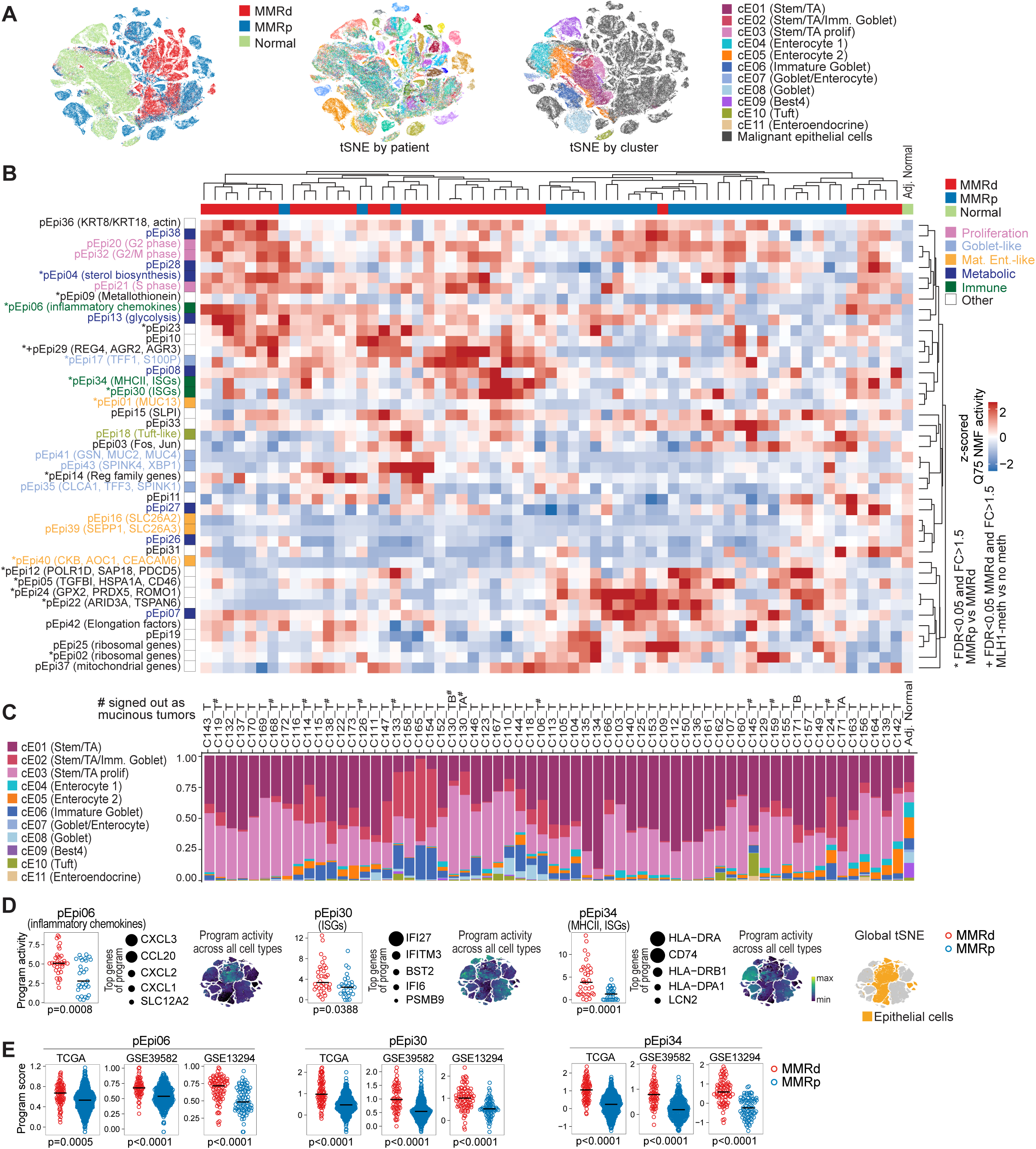
Transcriptional programs in malignant cells differ between MMRd and MMRp CRC. (**A**) tSNE of epithelial cells (left) with malignant epithelial cells clustering by patient (middle) and normal epithelial cells clustering by cell type (right). (**B**) Heatmap showing the 75th percentile of activities from the 43 malignant programs in malignant cells across CRC patient specimens (rows centered and z-scored), hierarchically clustered by average linkage. Gene program activity in normal epithelial cells is shown for comparison (rightmost column). Significant differences in MMRd vs MMRp are indicated by * (GLME (generalized linear mixed model), patient as random effects, MMR status as fixed effect, FDR<0.05). Significant difference between MLH1 promoter-methylated vs MLH1-non-methylated MMRd specimens is indicated with +. (**C**) Inferred cell-type composition of malignant cells in each tumor specimen, classified by supervised learning trained on non-malignant epithelial cells. Epithelial cell composition in normal tissue is shown for comparison (rightmost bar). Morphologically mucinous tumors are indicated with #. (**D**) Selected immune-related transcriptional programs with significantly different activity in MMRd vs MMRp CRC (GLME FDR<0.05). For each program the top genes are shown, circles indicate the relative weight of each gene in the program. tSNEs show program activities across all cell types. (**E**) Gene scores for programs in (D) in bulk RNA-seq of samples from TCGA-CRC (COADREAD), GSE39582, and GSE13294 patient specimens. Mann–Whitney–Wilcoxon test **** for p≤0.0001, *** for ≤0.001, **≤0.01, *≤0.05, ns for >0.05. See also Supplemental Figure 4 and **Supplemental Tables 4**,**6**.

Many programs were differentially active between malignant cells and normal epithelial cells. For example, mature enterocyte programs were reduced (Figure 4B **yellow**) and proliferation programs increased (Figure 4B **pink**) in malignant vs. normal epithelial cells, consistent with the vast majority of malignant cells being classified as stem/transit-amplifying (TA)-like cells (Figure 4C). Among the differentially active programs, 10 showed higher and 6 lower activity in MMRd compared to MMRp samples, a finding that we validated in three external datasets (Supplemental Figure 4A), along with similar grouping of programs across our cohort and in TCGA (Figure 4B; Supplemental Figure 4B).

In particular, three immune-related programs showed elevated activity between MMRd and MMRp malignant cells: an ISG (including IFNɣ targets; **Supplemental Table 6**) and MHC Class II gene program (pEpi34) was more active (3.4-fold) in MMRd than MMRp tumors; an ISG (Type I interferon targets; **Supplemental Table 6**) and MHC Class I gene program (pEpi30) was mildly elevated in MMRd vs. MMRp (1.6-fold; also with some activity in normal epithelial cells); and a neutrophil and immune-attracting chemokine program (*CXCL1,2,3 and CCL20*) (pEpi06) was higher in MMRd vs. MMRp tumors (1.6 fold) and in both tumor types compared to normal (Figure 4B **dark green, D,E; Supplemental Table 4**). Thus, malignant cells, especially in MMRd tumors, express immune-related gene programs that may mediate interactions with the immune system.

### Co-variation of program activities across patients predict multicellular immune hubs

We next hypothesized that some of the changes in gene programs within one cell type may be related to changes in another cell type, either because of a direct effect of one cell type on another, or because of a shared signal or neighborhood affecting both cell types in concert.

To find such networks of multi-cellular coordinated programs, we searched for program activities that are correlated across patient specimens (from hereon, ‘co-varying’ programs), analyzing MMRd and MMRp separately to better capture differences between the two immunologically disparate tumor types. We calculated pairwise correlations of program activities across each set of samples, using the 22 myeloid, 21 T/NK/ILC gene programs and either MMRd- or MMRp-derived malignant epithelial programs (EpiTd* and EpiTp*; Supplemental Figure 4E). Stromal cells were not included because the number of stromal cells per sample was insufficient for a co-variation analysis. Finally, we used graph-based clustering of programs (**Methods**) to identify 7 co-varying multi-cellular hubs in MMRd and 9 in MMRp samples (Figure 5A; Supplemental Figure 5A). These hubs consist of multiple programs expressed across the range of cell types, thus revealing multi-cellular interaction networks.

**Figure 5:**
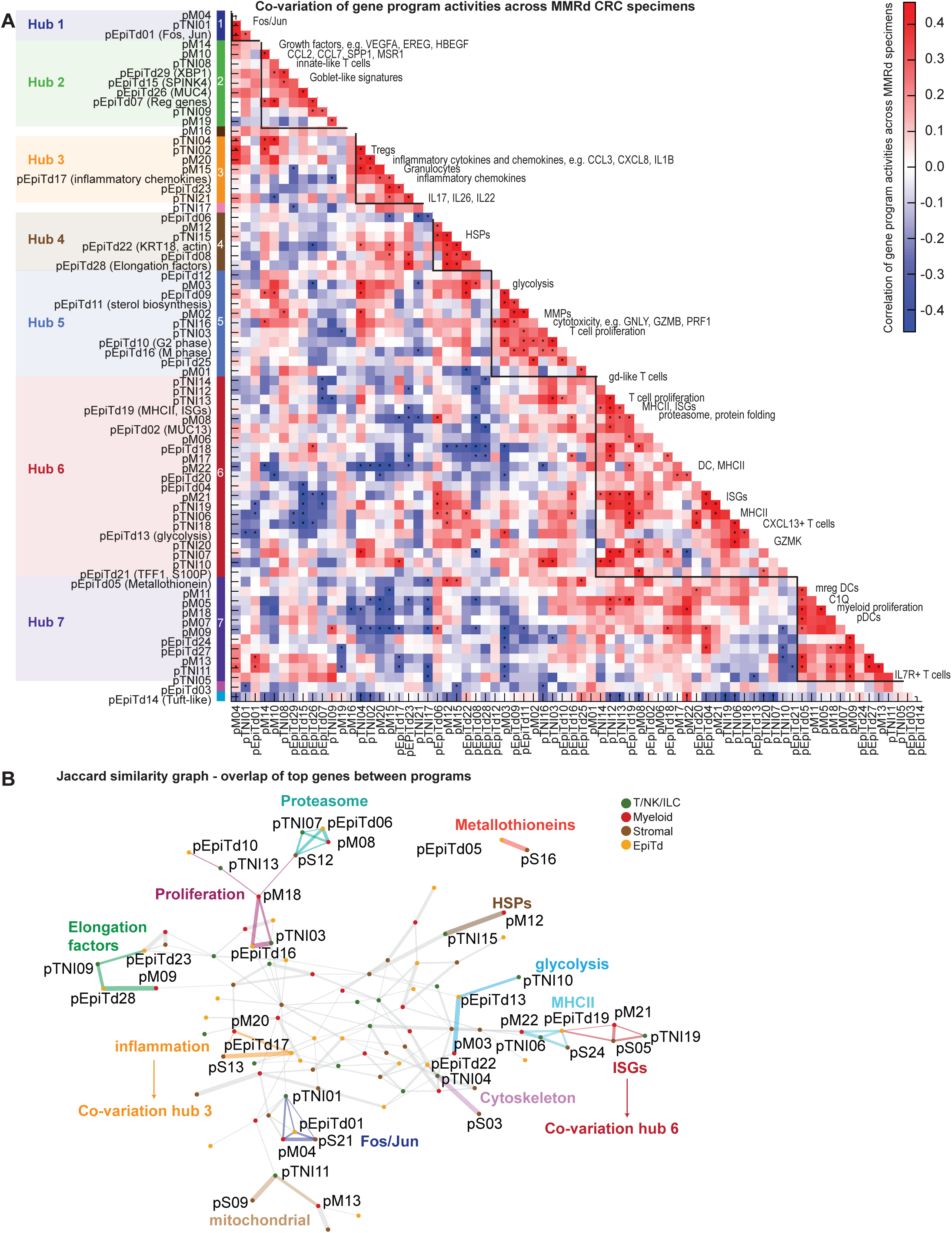
Discovery of multicellular interaction networks in MMRd CRC. (**A**) Heatmap showing pairwise correlation of gene program activities (‘co-variation score’) across MMRd specimens (**Methods**) using patient-level activities in T/NK/ILC, myeloid, and malignant compartment. Significance is determined using permutation of patient ID and is indicated with * (FDR<0.1). Densely connected modules (‘hubs’) are identified based on graph clustering of the significant correlation edges. (**B**) Similarity of transcriptional programs calculated based on the overlap of top weighted genes (Jaccard similarity) across T/NK/ILC, stromal, myeloid, and malignant cells (using the MMRd-derived malignant programs) visualized by nearest-neighbors graph. Edge thickness is proportional to program similarity. Edges from selected network neighborhoods are colored and annotated by function. See also Supplemental Figure 5 and **Supplemental Table 5**.

To identify programs that are similar to each other, and thus more likely to be triggered by a common mechanism, we computed the overlap of the top genes between programs. This analysis revealed immune, metabolic and other programs that were similar across cell types (Figure 5B, graph represents nodes as programs and edges as the Jaccard similarity; **Methods**). We note that co-varying programs (Figure 5A) need not be similar to each other (although they can be) and are often characterized by distinct top gene sets.

To study the interactions between malignant cells and immune cells, we focused on two MMRd-derived multicellular hubs (hub 3 and hub 6, Figure 5A) in which programs active in immune cells co-varied with immune-related programs active in malignant cells.

### Malignant cells, fibroblasts, monocytes, and neutrophils engage in inflammatory responses at the luminal surface of primary MMRd and MMRp tumors

Hub 3 featured inflammatory programs in malignant cells and monocytes that co-varied with a neutrophil program, all of which were highly active in both MMRd and MMRp tumors compared to normal tissue (Figure 6A, Supplemental Figure 6A). Treg and *IL17* T cell programs were also found in the hub. Hub 3 was active in MMRp samples (Supplemental Figure 5A, 6A), and its programs and their correlations were recapitulated in an external single cell cohort^18^ (Supplemental Figure 6A). Based on the similarity of myeloid, stromal and malignant inflammatory programs, which showed overlapping genes and shared transcription factor predictions, such as NF-κB, HIF1A, and CEPBP (Figure 6B), we also included stromal program pS13 (active in *GREM1*+ and *MMP3*+ CAFs; Figure 6C) in our analysis of hub 3.

**Figure 6:**
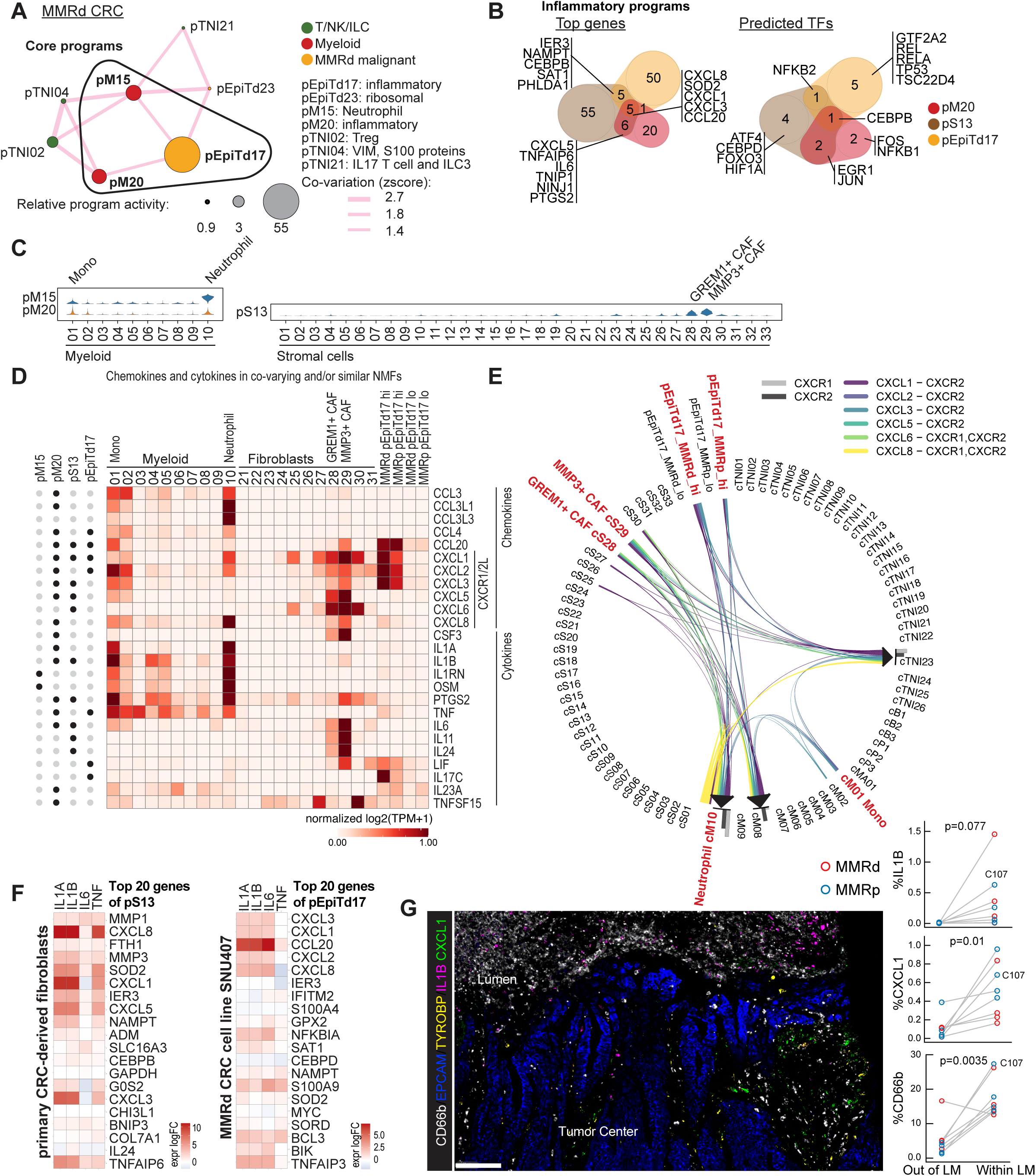
An inflammatory hub at the luminal surface of primary MMRd and MMRp tumors. (**A**) Inflammatory hub 3 is composed of significantly co-varying transcriptional programs across T/NK/ILC, myeloid, and malignant cells based on MMRd specimens. Node size is proportional to the log ratio of mean program activities in MMRd vs normal. Edge thickness is proportional to co-variation scores. Pink lines depict positive, blue lines negative correlations. (**B**) Venn diagrams showing the overlap of top weighted genes (left) and predicted transcription factors (right) for inflammatory gene programs in myeloid, stromal, and malignant compartment. (**C**) Violin plots showing program activities of pM15, pM20 across myeloid cell clusters and pS13 activity across stromal cell clusters. (**D**) Expression level of all chemokines and cytokines present in the top genes of the depicted NMF-based programs (indicated with black dot on the left) across the depicted clusters and malignant cells with high versus low pEpiTd17 program activity. Genes are normalized across all cell clusters in the data set (not only the clusters shown). (**E**) Interactions between CXCR1/2 and cognate chemokines. Clusters with high activity for the co-varying or similar inflammatory gene programs are marked in red. (**F**) Primary CRC-derived fibroblasts and SNU407 MMRd CRC cell line were stimulated with 10 ng/ml IL1A, IL1B, IL6, or TNF for 14h or left untreated. Transcriptional signature was determined by RNA-seq. Shown are log fold change (logFC) compared to unstimulated cells. Data are representative of two independent experiments each. (**G**) Multiplex RNA ISH/IF staining for neutrophils with *CD66b*-IF, epithelial cells with *EPCAM*-ISH, myeloid cells with *TYROBP*-ISH, the cytokine *IL1B*-ISH, and chemokine *CXCL1*-ISH. Representative image shows accumulations of neutrophils, *IL1B* and *CXCL1* signals at the malignant interface with the colonic lumen. Scale bar: 100um. Right, quantification of cell phenotypes in 8 CRC specimens shows that *IL1B*, *CXCL1*, and neutrophil (*CD66b*) signals are highly enriched in the luminal margin, defined as ≤ 360 um from the luminal border of the tumor. Paired two-tailed t-test. The patient not showing CD66b enrichment at the luminal margin is patient C110. Note 8 samples included at right because clinical paraffin block preparation required removal of luminal margin from one sample. See also Supplemental Figure 6 and **Supplemental Tables 5**,**6**.

To understand the communication pathways driving these malignant/immune/stroma cell interactions, we examined all chemokines and cytokines found within the top genes of the inflammatory and co-varying neutrophil programs (Figure 6D). This analysis suggested concerted attraction of *CXCR1/2*+ neutrophils by malignant cells, *GREM1*+ and *MMP3*+ CAFs, monocytes, and neutrophils expressing cognate chemokines (*CXCL1/2/3/5/6/8*) (Figure 6E). Indeed, the same chemokines were upregulated in CRC-derived fibroblasts and CRC malignant cells when we stimulated them *in vitro* with cytokines found in the hub 3 inflammatory monocyte and neutrophil programs, such as *IL1B* (Figure 6F). Malignant cells, CAFs, monocytes, and neutrophils thus appear to work in concert to recruit myeloid cells and amplify the recruitment of myeloid cells via inflammatory cytokines.

To localize this inflammatory hub within the tumor tissue, we stained MMRd and MMRp specimens for markers of neutrophils, myeloid cells, and malignant epithelial cells along with *IL1B* and *CXCL1* transcripts. 7 of 8 examined specimens showed significant accumulations of neutrophils along with *IL1B+* and *CXCL1*+ cells at the interface of the malignant cells with the colonic lumen (Figure 6G; Supplemental Figure 6B), particularly at sites with abundant necrosis. Although *CXCL1* was observed in malignant and myeloid cells, strong *CXCL1* signal was observed in cells that are neither myeloid nor epithelial. While these cells are likely to be the *MMP3*+ CAFs since they express the highest level of *CXCL1* by scRNA-seq (Figure 6D) and are mostly found at the luminal surface (Figure 3I), further imaging studies are needed to confirm this prediction. Taken together, given the localization of cells and molecules in this inflammatory hub (Figure 6G), and stromal remodeling (Figure 3I) at the luminal border, we suggest that damage at the luminal edge of primary colorectal cancers may contribute to positive inflammatory feedback loops that drive a myeloid and neutrophil-rich milieu in these tumors.

### A coordinated network of *CXCL13*+ T cells with myeloid and malignant cells

Hub 6 (Figure 5A, Figure 7A) was comprised of ISG/MHC-II gene programs expressed in both myeloid and malignant cells (likely induced by IFNɣ and driven by IRF/STAT transcription factors **Supplemental Table 6**, Figure 7B), which co-varied with *IFNG/MHC-II* and *CXCL13/PDCD1* T cell programs. These T cell programs include markers of activation and exhaustion (**Supplemental Table 2**) that are known to mark chronically stimulated tumor-reactive T cells^28, 50, 51^.

**Figure 7:**
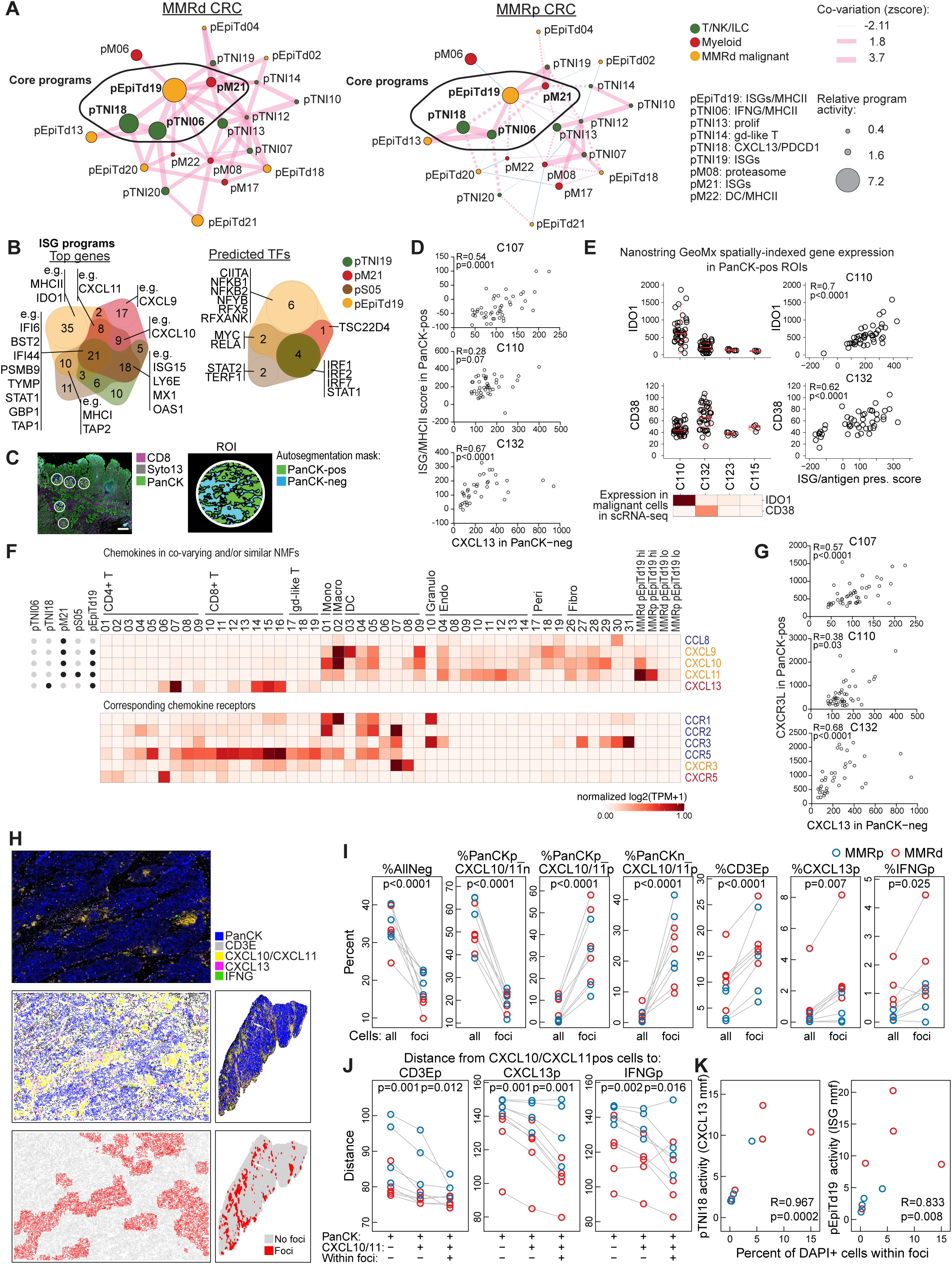
A coordinated network of *CXCL13*+ T cells with myeloid and malignant cells expressing ISGs. (**A**) Hub 6 is composed of significantly co-varying transcriptional programs across T/NK/ILC, myeloid, and malignant cells. Shown is hub 6 identified in MMRd specimens (left) and projected onto MMRp specimens (right). Node size is proportional to the log ratio of mean program activities in MMRd or MMRp vs normal. Edge thickness is proportional to co-variation scores. Pink lines depict positive, blue lines negative correlations. Non-significant edges are depicted as dotted lines. (**B**) Overlap of top weighted genes (left) and predicted transcription factors (right) for ISG programs in T/NK/ILC, myeloid, stromal, and malignant compartment. (**C**) Image shows a portion of the tissue from patient C110 with regions selected for spatially-indexed transcriptomics (GeoMx DSP CTA). ∼45 regions of interest (ROIs) per specimens were sampled and each ROI was auto-segmented based on the PanCK signal into PanCK-positive and -negative regions. Scale bar: 500 um. (**D**) Three CRC specimens with high *CXCL13* activity (namely C110, C132, and C107) were analyzed by spatially-indexed transcriptomics (GeoMx DSP CTA) as described in (C). *CXCL13* signal in PanCK-negative regions was correlated to an ISG/MHCII signal score in the paired PanCK-pos regions (Spearman correlation). ISG/MHCII score was calculated as described in Methods, based on: *HLA-DMA, HLA-DMB, HLA-DPA1, HLA-DQB1, PSMB10, PSMB8, PSMB9, TAP1, TAP2, TYMP, STAT1, CXCL10, CXCL11, GBP1, GBP2, GBP4*. (**E**) Quantification of NanoString GeoMx DSP CTA assay showing high *IDO1* expression in malignant cells of patient C110, and high *CD38* expression in malignant cells of C132, consistent with scRNA-seq data (heatmap below left scatter plots, log2TPM). Right: Scatter plots and Spearman correlation between *IDO1* (top) or *CD38* (bottom) expression and the ISG scores (as calculated in D) in malignant cells of the respective patients. (**F**) All chemokines present in the top genes of the depicted NMF-based programs (indicated with black dot on the left) as expressed in the depicted clusters and malignant cells with high versus low pEpiTd19 program activity. Genes are normalized across all cells clusters in the data set (not only the clusters shown). (**G**) GeoMx DSP CTA assay as in (D) shows that *CXCL13* signal in PanCK-negative regions is correlated to the expression of CXCR3 ligands (determined as sum of *CXCL9*, *CXCL10*, *CXCL11*) in the paired PanCK-positive regions (Spearman correlation). (**H**) Multiplex RNA ISH / IF staining for PanCK-IF, *CD3E*-ISH, *CXCL10/CXCL11*-ISH, *CXCL13*-ISH, and *IFNG*-ISH was performed on 9 tumor tissue slides from different donors (MMRd n=5: C110, C123, C132, C139, C144; MMRp n=4: C103, C112, C126, C107) and cells were phenotyped using the Halo software. An image section from C123 is shown (top), a computational rendering of the same section (middle left) and the full slide (middle right). Cells were characterized by a 100µm neighborhood and clustered by their neighborhood features to identify two clusters (‘foci’ and ‘no foci’). Scale bar: 100um. (**I**) Based on the approach as outlined in (H), percentages of the indicated phenotype (p: positive; n: negative) among either all DAPI+ cells or the DAPI+ cells within the foci were calculated. *CXCL10/CXCL11*p, *CD3E*p, *CXCL13*p, and *IFNG*p cells are significantly enriched in foci. (**J**) Distances were calculated from *CXCL10*/*CXCL11*-positive cells to the indicated phenotypes (mean distance across 100um neighborhoods) outside or inside the foci. If a phenotype was not observed in the 100um neighborhood, the distance was set to 150um. (**K**) Percentage of cells within foci (among all DAPI+ cells) was correlated to scRNA-seq-based pTNI18 and pEpiTd19 activities from the respective specimens (Spearman correlation). See also Supplemental Figure 7 and **Supplemental Tables 5,6,7**.

Importantly, we did not derive this hub in an MMRp-specific analysis (Supplemental Figure 5) and observed weaker activities of the core programs and reduced connectivity (*e.g,* the link between malignant pEpiTd19 and T cell pTNI18 programs is lost) of the network when we projected the network onto MMRp tumors in our dataset and an external scRNA-seq dataset (Figure 7A, Supplemental Figure 7A), consistent with the weaker immunogenicity of MMRp tumors.

To validate the co-activity of ISG/MHC-II malignant and *CXCL13* T cell programs, we performed spatially-indexed transcript profiling (GeoMx® Digital Spatial Profiling, **Methods**) of tissue sections from three CRC tumors that showed high *CXCL13* T cell program activity in matching scRNA-seq data. We profiled 45 regions of interest (ROI) per tumor section, and further segmented each region into epithelial vs non-epithelial areas (Figure 7C). We observed a positive correlation between ISG expression in malignant epithelial areas and *CXCL13* expression in adjacent non-epithelial areas across all regions per tumor (Figure 7D), further supporting potential interactions between malignant and T cells in this hub.

Consistent with the presence of exhausted T cells in this hub, the malignant ISG/MHC-II program also featured inhibitory molecules, including transcripts encoding the enzymes *IDO1* and *CD38* that were included as probes in the GeoMx® analysis. *IDO1* and *CD38* expression in the malignant ROI of 4 patients was comparable to expression measured by scRNA-seq for the same patients. Moreover, *IDO1* or *CD38* expression was spatially correlated with ISG scores (Figure 7E) in patients with high scRNA-seq-derived expression of these two genes and the *CXCL13* T cell program. These results show that negative feedback mechanisms are part of the hub’s function, and that the inhibitory feedback system is regulated by patient-specific and region-specific factors in each tumor.

### *CXCL13*+ T cells are localized within foci of *CXCL10/CXCL11*-expressing cells throughout the tumor

Given spatially correlated expression of ISGs in malignant cells with *CXCL13* in non-malignant regions (Figure 7D), we hypothesized that T cells would be spatially organized around cells expressing T cell attracting chemokines. We examined all chemokines in the hub 6 gene programs, and found that myeloid, malignant and stromal ISG programs included the chemokines *CXCL9*, *CXCL10*, and *CXCL11* (Figure 7F), and that their cognate receptor *CXCR3* was upregulated in activated T cells and certain DC subsets (Figure 7F). Using our spatially-indexed transcriptomic dataset of three highly T cell infiltrated samples (patients C107, C110, C132), we validated this observation by finding that C*XCL13* expression in non-epithelial cells was associated with *CXCR3* ligand expression in the malignant cells of the same ROI (Figure 7G).

To further validate this spatial association at single cell resolution, we performed whole section staining of nine CRC specimens from our scRNA-seq cohort (Figure 7H-K, **Supplemental Table 7**). We found that *CXCL10/CXCL11*-positive cells were clustered into large foci enriched for cells expressing *CXCL13* and/or *IFNG*, as well as *CD3E*+ T cells (Figure 7H, I, Supplemental Figure 7B**, Methods**). Interestingly, foci in specimens with high (3 MMRd and 1 MMRp) versus low (2 MMRd and 3 MMRp) *CXCL13*+ T cell program activity tended to show *CXCL10/CXCL11* expression in malignant cells versus non-malignant cells, respectively (Supplemental Figure 7B,C), though additional studies are needed to confirm this observation.

Across all samples, *CXCL10/CXCL11*+ malignant epithelial cells were on average closer to *CD3E*+, *CXCL13*+, and *IFNG*+ cells than their *CXCL10/CXCL11*-negative counterparts, and these distances were especially small within foci (Figure 7J). Lastly, specimens with greater scRNA-seq-derived activity of pTNI18 (*CXCL13* program) and pEpiTd19 (ISG program) had more cells participating in *CXCL10/CXCL11* foci (Figure 7K, Supplemental Figure 7B). Our findings thus reveal spatially organized foci consisting of activated *IFNG*+ and *CXCL13*+ T cells together with *CXCL10/CXCL11+* myeloid and malignant cells, providing evidence that a positive feedback loop – by which T cell-derived IFNɣ induces expression of CXCR3 ligands to attract more T cells – may be critical in the formation of these immune cell hotspots within tumors.

## Discussion

Tumors are very heterogeneous as a result of genetic evolution and epigenetic variation, but the immune cells infiltrating tumors are less plastic and exhibit a more limited set of behaviors. To identify recurring cell-cell interactions that contribute to the immune response in colorectal cancer, we considered not only cell composition, but also cell intrinsic expression programs, and the dependencies between programs of different cells within and across tumors. This allowed us to define hubs of interacting cells based on co-varying cellular programs, followed by spatial profiling to ask how components of these hubs are spatially organized in the tumor. The resulting set of validated and inferred hubs provides a map of coordinated multi-cellular responses in both MMRd and MMRp tumors.

Our study shows that T cells are organized in structured cell neighborhoods within human tumors. We found that hub 6 consisted of interacting malignant cells, T cells and other immune cells localized deep within the tumor. The formation of these hotspots likely depends on a positive feedback loop in which T cell-expressed *IFNG* drives the induction of *CXCR3* chemokines (as part of the ISG response) that then attract more T cells and other cells. Supporting this notion, recent studies showed that expression of *CXCR3* chemokines in myeloid cells is required for inducing anti-tumor T cell responses following checkpoint inhibitor treatment in mice^52, 53^. Furthermore, several studies have linked the *CXCR3* chemokine system to T cell entry into tissues, including CD8+ T cell recruitment in melanoma^54^, viral infection in which *CXCR3* ligands are induced by CD4+ T cell-derived IFNɣ^55^, and vaccination in which topical CXCL9 and CXCL10 administration recruited activated T cells into epithelial tissue, even in the absence of antigen^56^. In humans, an IFNɣ-induced signature^29, 57^, which overlaps with the genes we observed in the programs of hub 6, was associated with favorable response to PD-1 blockade in multiple human tumor types. In contrast to the positive feedback loop, persistent ISG hubs in tumors may drive immunosuppression due to a negative feedback loop that upregulates co-inhibitory factors such as *PD1/PDL1*, *Lag3/MHCII*, *Tim3/LGALS9*, and *IDO1*. Indeed, mechanistic work in the B16 melanoma mouse model suggests that IFNɣ can drive a multigenic resistance program^58^. Whether the positive or negative feedback is dominant at a particular location or time will be important to determine in different tumor types and treatments.

Another important question is whether these multicellular immune formations are similar to previously observed lymphoid structures or aggregates in tissues. Tertiary lymphoid structures (TLS)^59^ contain germinal center B cells, and have been associated with high T cell activity, favorable prognosis and effective response to immunotherapy^59–63^. In contrast to TLS which are often found below the invasive margin of tumors^16^, hub 6 was found in the tumor center and does not harbor germinal centers, and thus is not likely to be a mature TLS. Consistently, we observed a depletion of B cells in our samples relative to normal colon. A few studies observed aggregates that are not likely to be TLS. In an early study of melanoma immunity, staining for IFNɣ, T cells and PD-L1 showed their spatial proximity in tumors^64^. Another group observed aggregates of stem-cell-like CD8+ T cells with MHCII+ cells, which were associated with less progressive kidney cancer in human^65^. A third study showed that vaccination of mice induced an IFNG/CXCR3-dependent spatial hub of T cells and myeloid cells expressing CXCL10. This hub formed around the vasculature and facilitated entry of circulating T cells into the tissue^66^, thus providing a platform for frequent encounters of T cells with other cells to coordinate immune responses.

The other hub was centered around an inflammatory positive feedback loop between inflammatory CAFs, monocytes, and neutrophils driven by cytokines such as *IL1B* and the *CXCR1/2* chemokine system and located at the luminal surface. The luminal surface of colonic tumors has an abnormal epithelial lining and the tumor mass protrudes into the gut lumen where it can suffer abrasive injury from colonic contents. Tissue damage could lead to entry of microbial ligands or release of immunostimulatory ligands from dead cells, resulting in inflammation and formation of granulation tissue. We also observed dilated blood vessels at the luminal surface, consistent with previous studies in CRC^67^. We found highly inflammatory fibroblasts expressing MMPs around the dilated vessels; these MMPs are known to contribute to tumor angiogenesis^68^. The inflammatory fibroblast program also included components of the prostaglandin pathway – *PTGS2* (encodes COX2) and *PTGES* (encodes prostaglandin E synthase) – that can promote CRC growth and suppress anti-tumor immune response^69^. Indeed, inhibition of the prostaglandin pathway is a major therapeutic focus in the prevention and treatment of CRC^69^. The inflammatory hub furthermore featured the Treg program and a T cell program including IL17, both of which were implicated in the suppression of anti-tumor responses and promotion of tumor growth. IL17 has been shown to promote angiogenesis and tumor expansion in murine models^70–72^, including through CAF activation and recruitment of granulocytes that can support tumor growth^71, 72^.

Our study provides a rich dataset of cellular states, gene programs and their transformations in tumors (such as the profound changes observed in stromal cells) across a relatively large cohort of patients with colorectal cancer. Our predictions of several multicellular hubs based on co-variation of gene expression programs, and subsequent spatial localization of two major immune-malignant hubs, organizes a large set of cell states and programs into a smaller number of coordinated networks of cells and processes. Understanding the molecular mechanisms underlying these hubs, and studying their temporal and spatial regulation upon treatment will be critical for advancing cancer therapy.

## Acknowledgements

We would like to thank the Broad Genomics Platform, Broad Flow Cytometry Facility, and Pathology and Surgery Departments at MGH and BWH. We thank the members of the Villani, Regev and Hacohen labs for advice and support throughout the project This work was made possible by the generous support of the Evergrande Center for Immunologic Diseases at Brigham and Women’s Hospital and Harvard Medical School (A.C.A., A.M.M.), Klarman Cell Observatory (O.R., A.R.), HHMI (A.R); NIH/NCI R01 CA208756 (N.H.), and the Arthur, Sandra and Sarah Irving Fund for Gastrointestinal Immuno-Oncology (N.H.). The project was also funded in part with Federal funds from the National Cancer Institute, National Institutes of Health, Task Order No. HHSN261100039 under Contract No. HHSN261201500003I and is part of the NIH HTAN and HTAPP consortium. The content of this publication does not necessarily reflect the views or policies of the Department of Health and Human Services, nor does mention of trade names, commercial products, or organizations imply endorsement by the U.S. Government. We also thank the following funding sources: Research fellowship of the German Research Foundation (DFG), Stand Up to Cancer Peggy Prescott Early Career Scientist Award PA-6146, Stand Up to Cancer Phillip A. Sharp Award SU2C-AACR-PS-32 and BroadIgnite (K.P.); NIH/NCI T32CA207021 (J.H.C.) The Doris Duke Charitable Foundation, The Pancreatic Cancer Action Network, NIH-NCI K08 CA218420-02, P50 CA127003, U01 CA224146 (A.J.A); Stand Up to Cancer Colorectal Dream Team Translational Research Grant (SU2C-AACR-DT22-17) (R.B.C); NIH grant R35 CA197735 (S.O.); K08CA222663, U54CA225088, Burroughs Wellcome Fund Career Award for Medical Scientists, CUMC Louis V. Gerstner, Jr. Scholars Program, CUMC Velocity Fellow Program (B.I.); NIH/NCI R01CA205406, DOD CA160344, Project P Fund (K.N.); U54 CA224068 (R.C., N.H); Stand Up to Cancer Colorectal Cancer Dream Team Translational Research Grant SU2C-AACR-DT22-17 (R.C., N.H), administered by the American Association for Cancer Research, a scientific partner of SU2C, Conquer Cancer Foundation of ASCO Career Development Award (M.G.); U2C CA233195 (B.E.J.). Nir Hacohen, PhD, is the David P. Ryan, MD Endowed Chair in Cancer Research, funded by a Gift from Arthur, Sandra and Sarah Irving.

## Author Contributions

Conceptualization: K.P., M.H., J.H.C., S.S., B.I., A.J.A., A.C.A., O.R., A.R., N.H.; Methodology: K.P., M.H., J.H.C., S.S., J.D.P., V.J., M.H., M.B.; Software: M.H., S.S., A.B., K.X.. C.S., C.L., L.T.N.; Formal analysis: K.P., M.H., J.H.C., S.S., J.D.P., V.J., A.B., K.X., S.C., D.Z., D.L., J.R., O.C., L.T.N.; Investigation: K.P., J.H.C., J.D.P., V.J., D.D., W.G., D.Z., K.R., L.N., J.W., I.W., M.K., J.W., M.J.S, J.Y., B.V., A.M., A.K.; Resources: D.Z., J.R., K.R., M.H., T.C., R.B., N.M., J.I., H.K., D.L.B., A.S., J.L.H., S.O., K.N.; Data Curation: J.H.C., L.D., S.D., D.F., T.S., K.N., M.G., A.J.A.; Writing - Original Draft: K.P., M.H., J.H.C., S.S., N.H.; Writing - Review & Editing: K.P., M.H., J.H.C., S.S., J.D.P., S.C., S.O., B.E.J., K.N., M.G., L.T.N., A.J.A., A.C.A., O.R., A.R., N.H.; Supervision: K.P., M.H., J.H.C., A.R., S.V., C.B., M.G., L.T.N., G.M.B., A.J.A., A.C.A., O.R., A.R., N.H.; Project administration: D.D., T.D., T.M., M.G-R., A.A., L.B., L.D., S.D., S.P., E.H., J.M., D.F., T.S., E.Z.T., J.J-V.; Funding acquisition: K.P., B.I., B.E.J., R.B.C., A.H.S, V.K.K., K.N., G.M.B., A.J.A., A.C.A., O.R., A.R., N.H.

## Supplemental Figure Legends

**Supplemental Figure 1:**
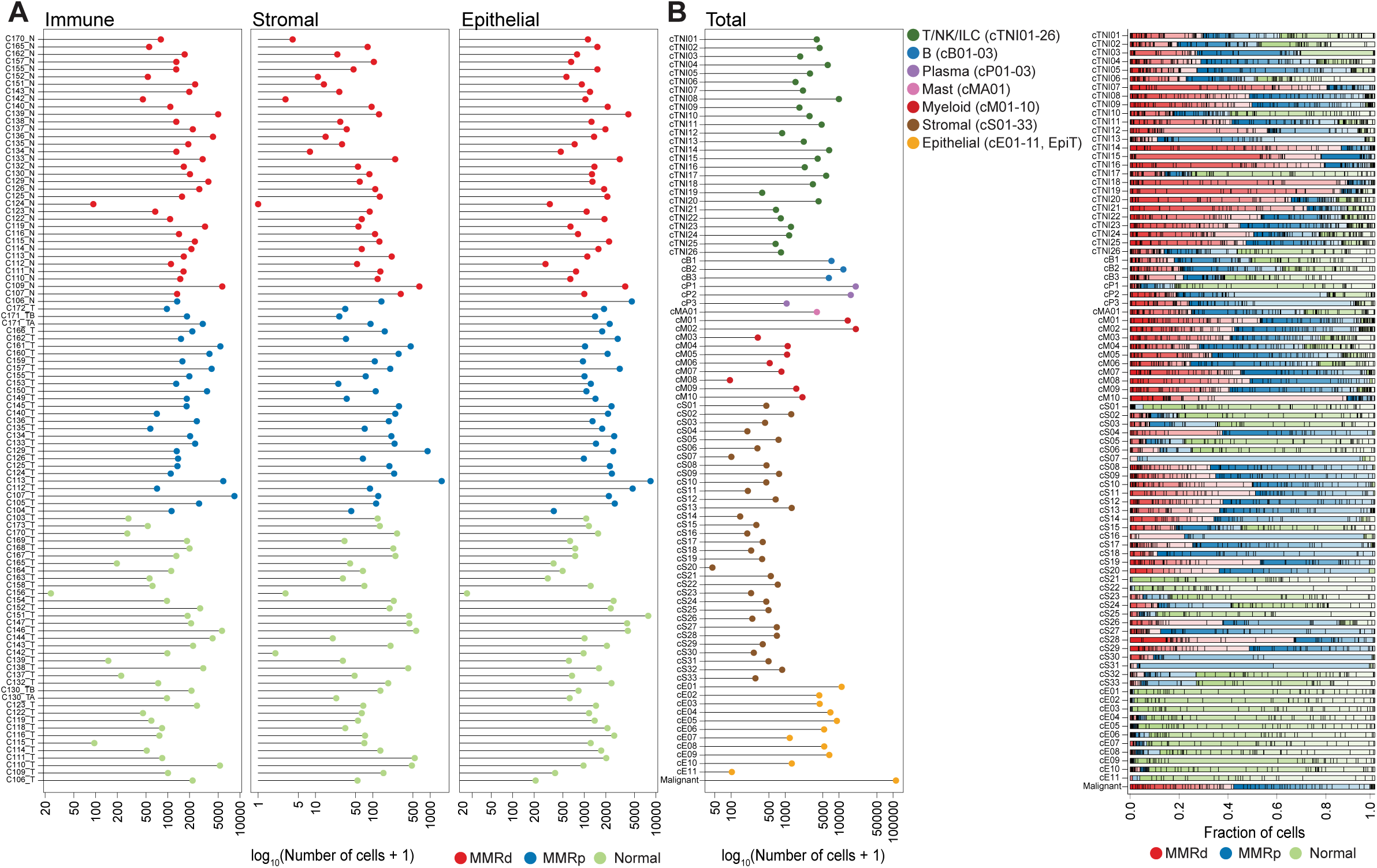
Cellular composition of scRNA-seq data set. (**A**) Number of cells in immune (T/NK/ILC, B, Plasma, Mast, Myeloid), stromal (Endothelial cells, Pericytes, Fibroblasts, Smooth Muscle cells, Schwann cells), and epithelial (malignant in tumor and non-malignant in normal specimens) compartment per specimen. (**B**) Number of cells per cluster (left) and fraction of cells from MMRd, MMRp, and normal specimens (right) within each cluster. Each specimen is indicated by a different color shade and separated by a vertical black line.

**Supplemental Figure 2:**
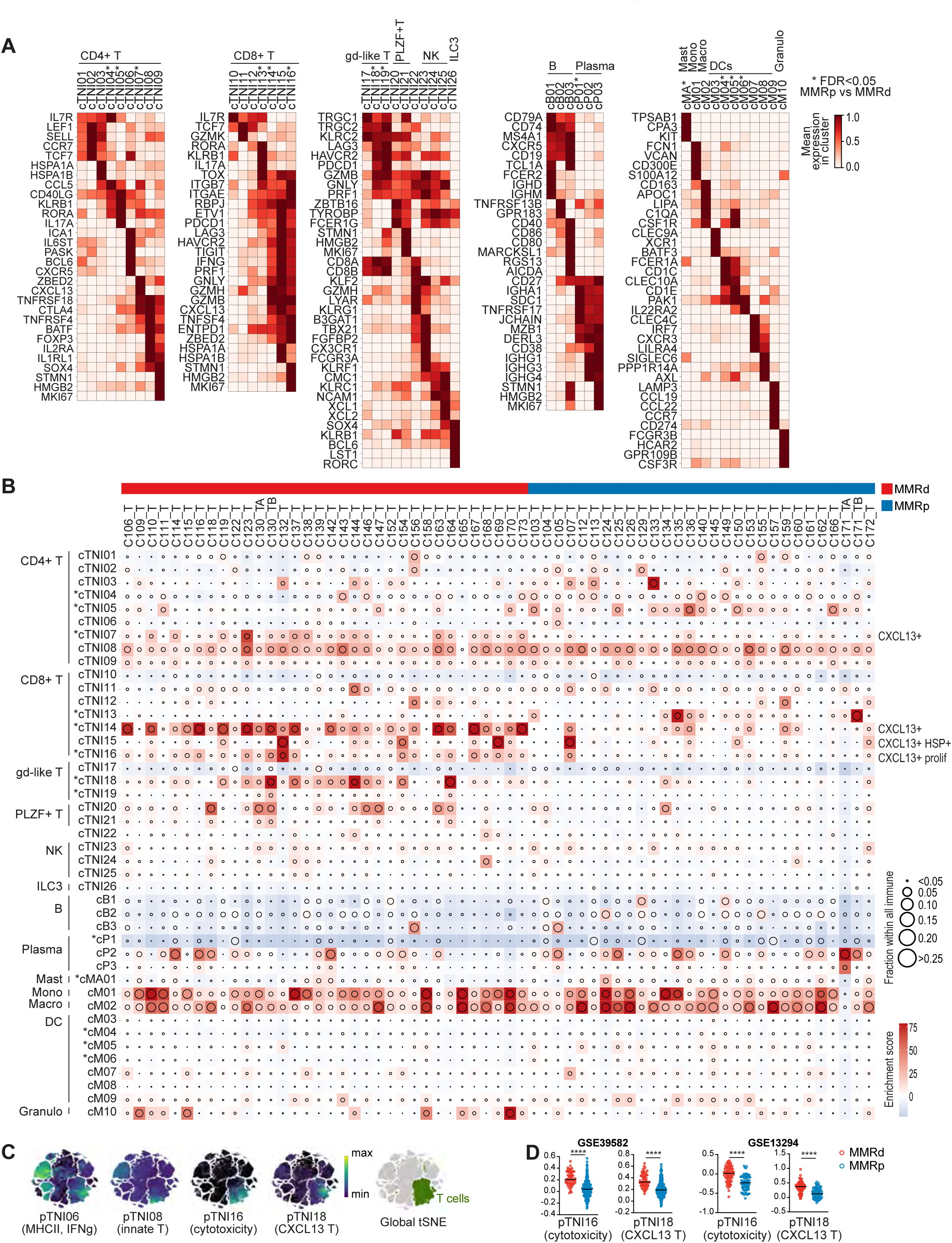
The immune compartment in MMRd and MMRp colorectal cancer. (**A**) Heatmaps showing selected unbiased and well established marker genes for immune clusters as mean expression in normalized log2(TPM+1). A comprehensive list of DEGs for each cluster can be found in **Supplemental Table 2**. (**B**) Changes in immune cell clusters in MMRp and MMRd tumors relative to adjacent normal tissue, showing frequency of immune cells (dot size) or enrichment/depletion (Pearson residual, colored squares). Clusters with differences in frequency between MMRp and MMRd tumors with FDR<0.05 are marked with *. (**C**) tSNE showing pTNI06, pTNI08, pTNI16, and pTNI18 program activities (**D**) Gene signature score for pTNI16 and pTNI18 in MMRd and MMRp CRC in bulk RNA-seq from GSE39582 and GSE13294 patient specimens (Mann–Whitney–Wilcoxon test with ns for p>0.05, * for p≤ 0.05, ** for p≤ 0.01, *** for p≤ 0.001, **** for p≤ 0.0001).

**Supplemental Figure 3:**
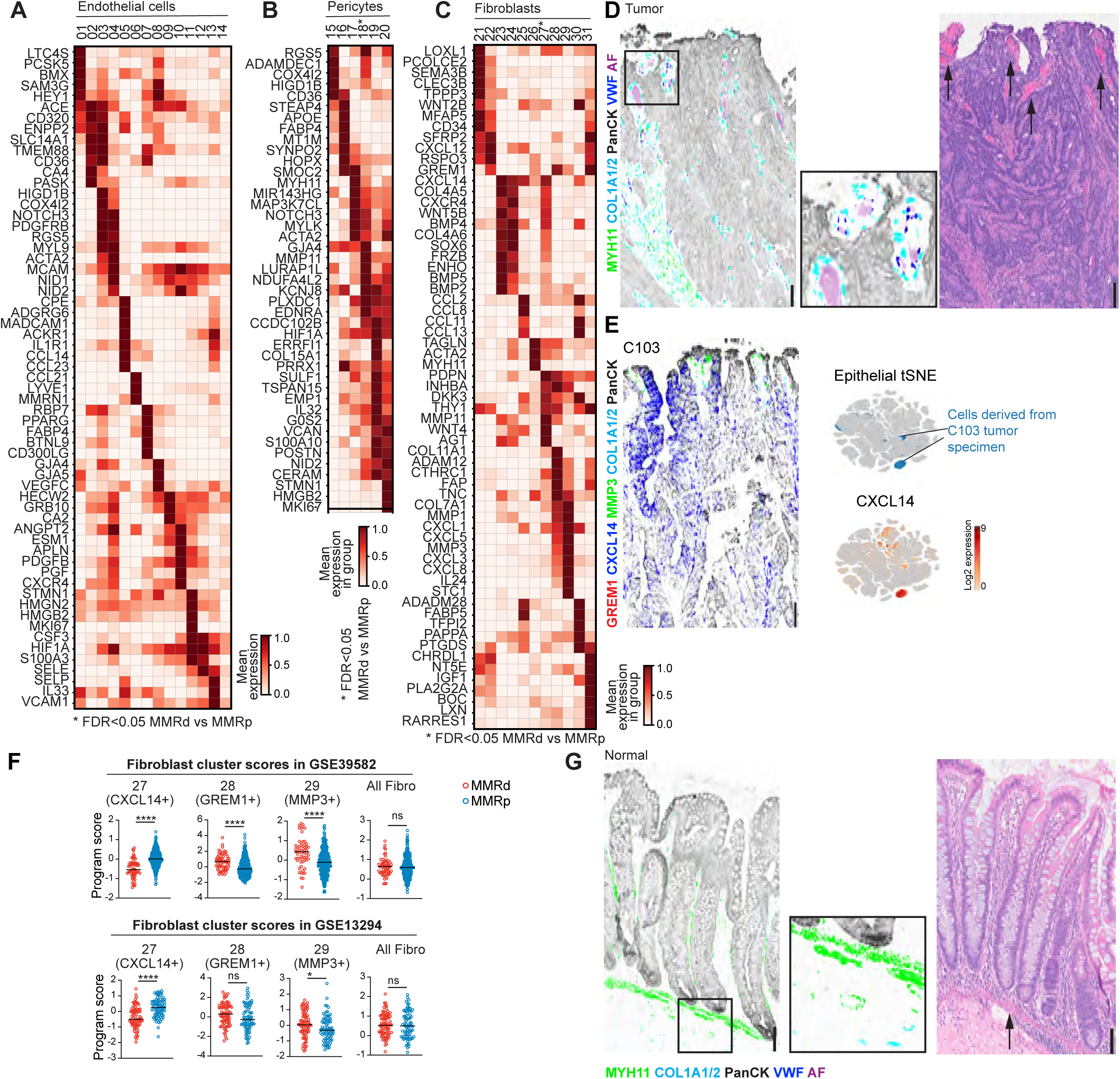
Stromal remodelling in MMRd and MMRp CRC. (**A**) Heatmap showing selected unbiased and well established marker genes for endothelial cell clusters as mean expression in normalized log2(TPM+1). A comprehensive list of DEGs for each cluster can be found in **Supplemental Table 3**. (**B**) As (A) for pericyte clusters. (**C**) As (A) for fibroblast clusters. (**D**) Serial section of the same area as in Figure 3I and 3L stained by multiplex RNA ISH/IF for smooth muscle marker *MYH11*-ISH, fibroblast marker *COL1A1/2*-ISH, epithelial marker PanCK-IF, and endothelial marker VWF-ISH (left image). The *MMP3*+ CAFs surround *VWF*+ endothelial cells enclosing autofluorescent (AF) red blood cells. H&E image (right) with dilated blood vessels (bright pink, marked with arrows). Scale bars: 100um. (**E**) Representative multiplex RNA ISH/IF image of patient C103 showing *CXCL14*-ISH expression by both epithelial lining fibroblasts and by the malignant epithelial cells. Scale bar: 100um. tSNE shows *CXCL14* expression in the malignant cells of patient C103 by scRNA-seq. (**F**) Gene signatures of *CXCL14*+ CAFs, *GREM1*+ CAFs, *MMP3*+ CAFs, and all fibroblasts in MMRd and MMRp bulk RNA-seq of patient specimens in GSE39582 and GSE13294. Mann–Whitney–Wilcoxon test **** for p≤0.0001, *** for ≤0.001, **≤0.01, *≤0.05, ns for >0.05. (**G**) Representative multiplex RNA ISH/IF (as in D) image of *MYH11*+ *COL1A/2*-negative muscularis mucosa below the base of the crypt in non-neoplastic colon (left). H&E image (right) of the same region with arrow pointing to muscularis mucosa. (**H**) Consecutive section to Figure 3L stained with multiplex RNA ISH/IF showing very little *INHBA*-ISH in non-neoplastic colon (left and lower-middle inset), and substantial *INHBA* staining in CAFs (upper-middle inset and right image).

**Supplemental Figure 4:**
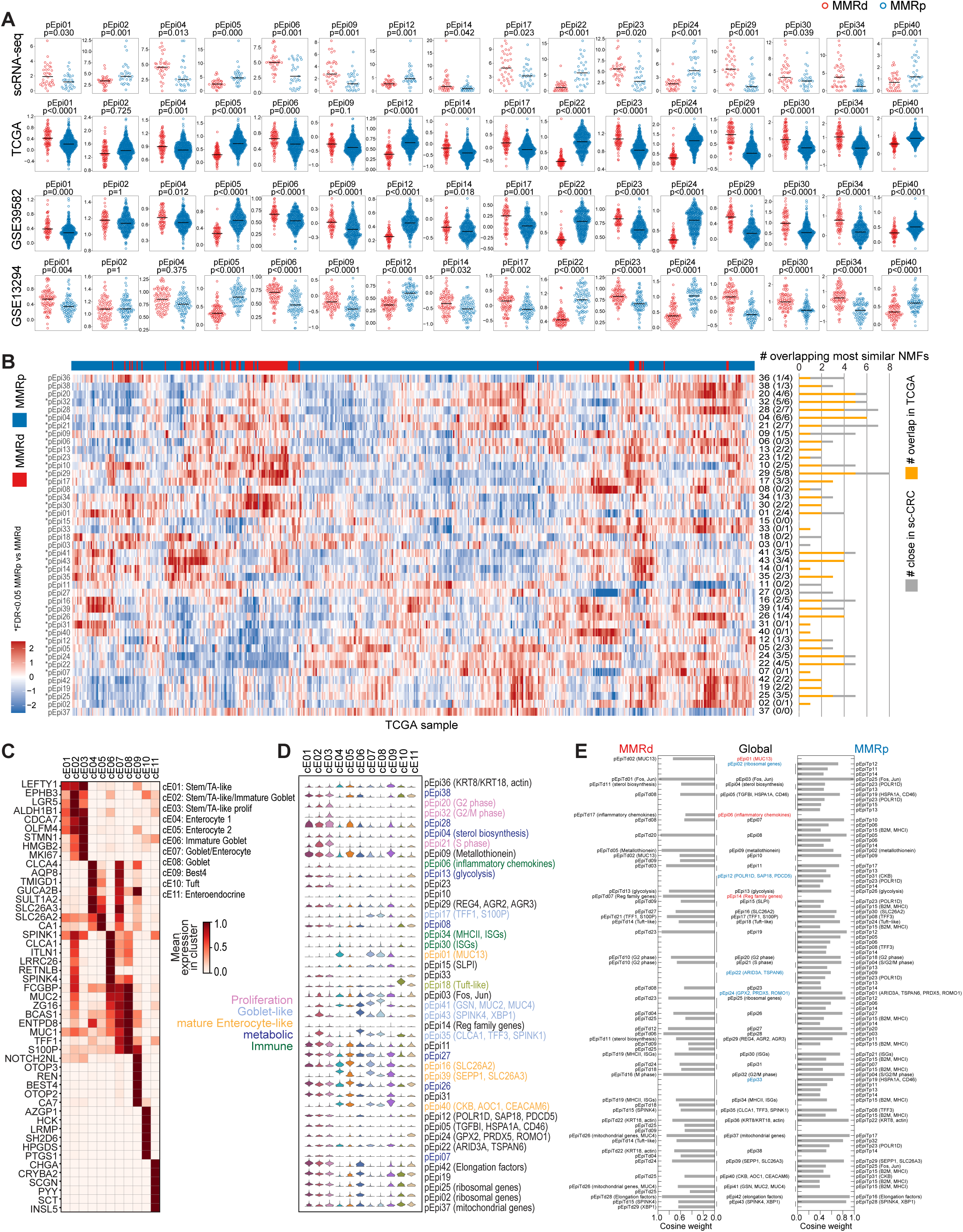
The epithelial compartment in adjacent normal colon tissue, MMRd and MMRp CRC. (**A**) Epithelial programs with significantly differential activities between MMRd and MMRp tumors in the scRNA-seq data set (GLME (generalized linear mixed model) FDR<0.05 and >1.5-fold difference between means) scored in bulk RNA-seq from three external cohorts. (**B**) Gene signature for the 43 epithelial programs in bulk RNA-seq from TCGA-CRC (COADREAD) patient specimens. Rows are ordered as in Figure 4B, columns are clustered. Significant MMRd versus MMRp differences are marked with * (Wilcoxon, two-sided with family-wise error rate corrected P≤0.05). Bar to the right of the heatmap shows the number of most closely correlated programs (≥90th percentile of correlations) based on program activities within scRNA-seq data (yellow+grey) and number of those most closely correlated programs that are preserved in TCGA (yellow). (**C**) Heatmap shows selected unbiased and well established marker genes for normal epithelial cell clusters. A comprehensive list of DEGs for each cluster can be found in **Supplemental Table 4**. (**D**) Transcriptional activities of epithelial programs within normal epithelial cell clusters. (**E**) Similarity between epithelial gene programs and MMRd- and MMRp-derived gene programs based on cosine weight. Programs that only had close matches in MMRd are marked in red, programs that only had close matches in MMRp are marked in blue. See also **Supplemental Table 4**.

**Supplemental Figure 5:**
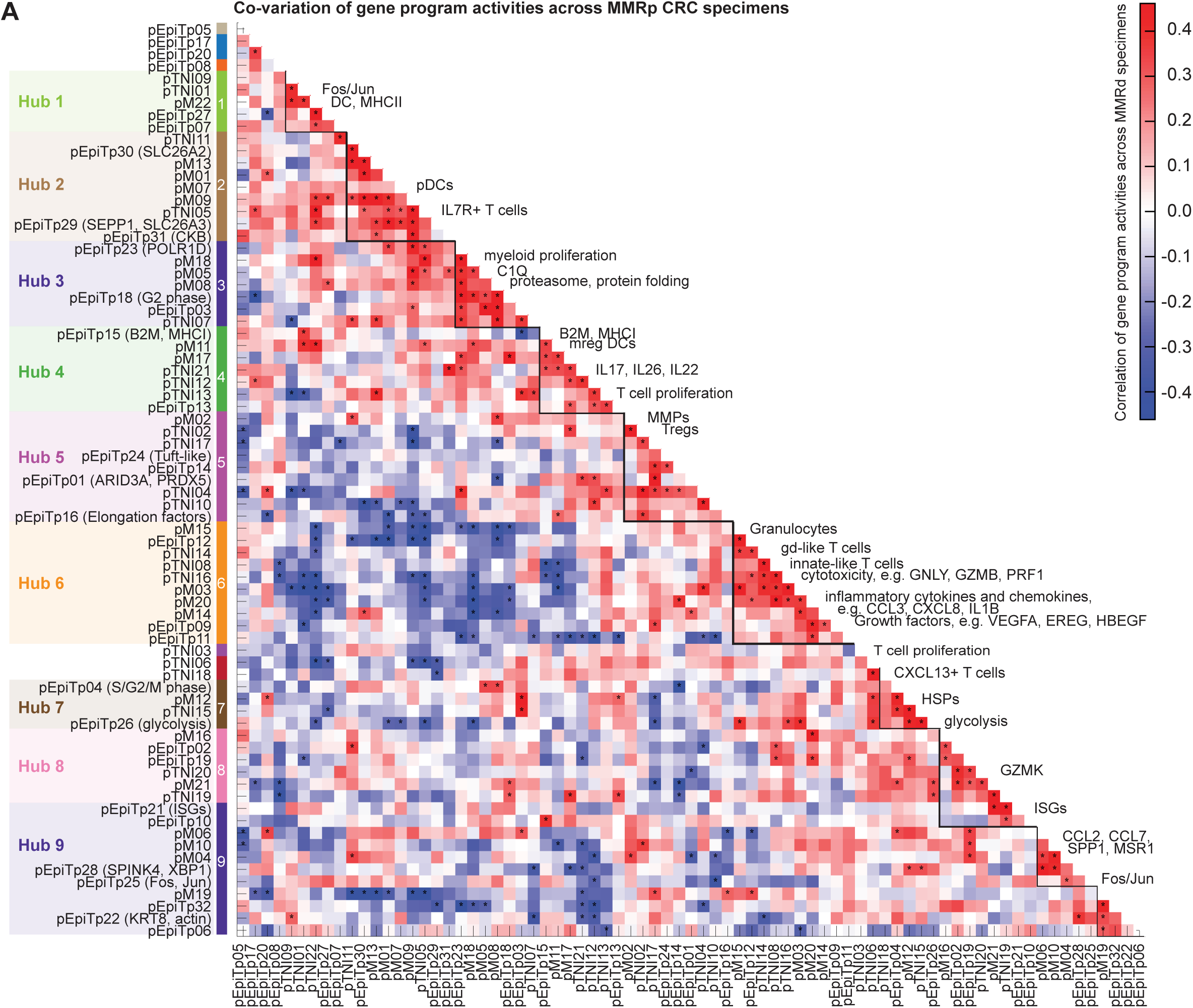
Discovery of multicellular interaction networks in MMRp CRC. (**A**) Heatmap showing pairwise correlation of gene program activities (‘co-variation score’) across MMRp specimens (**Methods**) using patient-level activities in T/NK/ILC, myeloid, and malignant compartment. Significance is determined using permutation of patient IDs and is indicated with * (FDR<0.1). Densely connected modules (‘hubs’) are identified based on graph clustering of the significantly correlated edges.

**Supplemental Figure 6:**
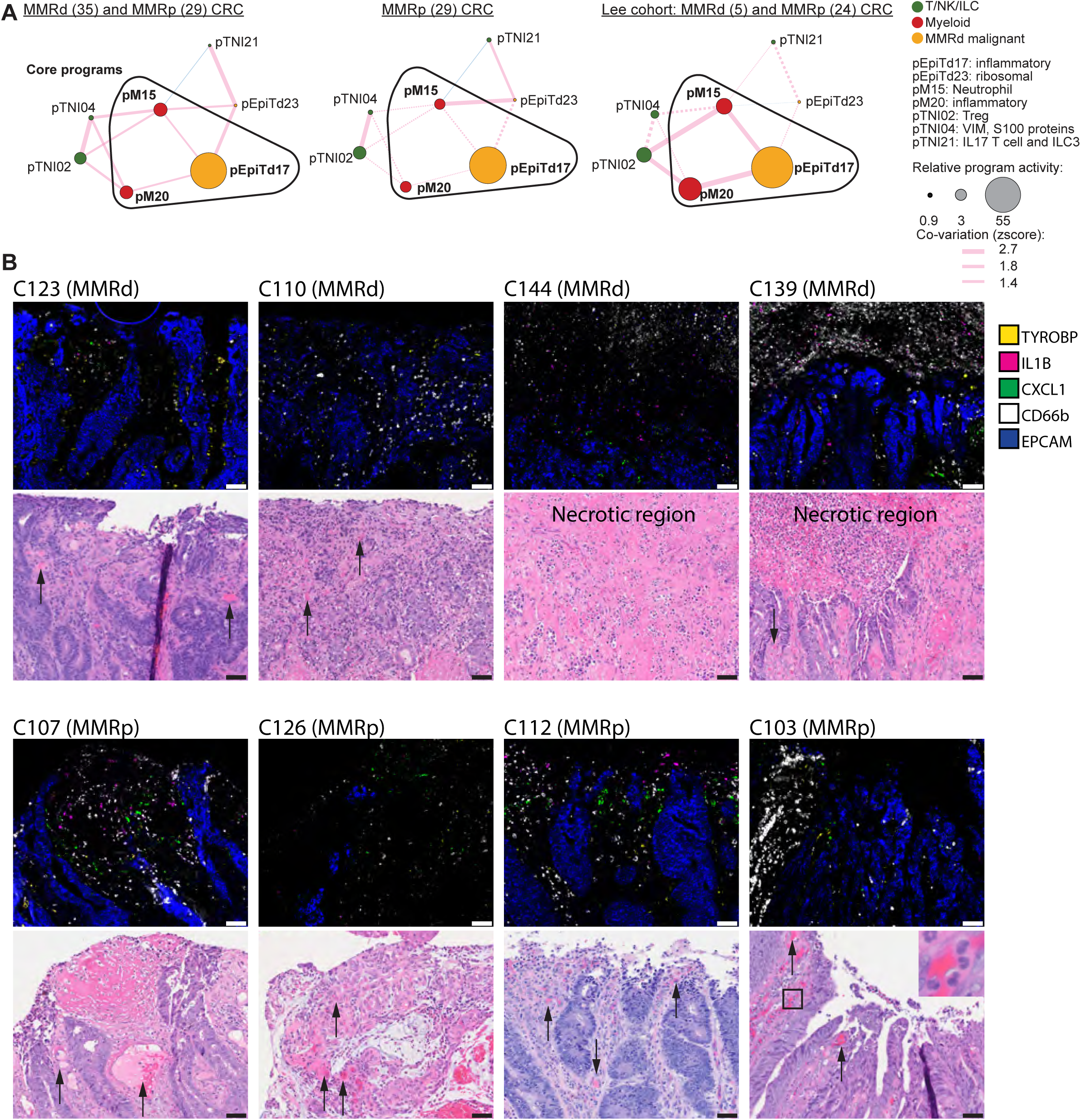
An inflammatory hub at the luminal surface of primary MMRd and MMRp tumors. (**A**) Inflammatory hub 3 as discovered in MMRd projected onto all MMRd and MMRp specimens from our scRNA-seq cohort (left; n=35 MMRd, n=29 MMRp), MMRp specimens (middle) or ^18^ (right; n=5 MMRd, n=24 MMRp). Node size is proportional to the log ratio of mean program activities in MMRd or MMRp vs normal. Edge thickness is proportional to co-variation scores. Pink lines depict positive, blue lines negative correlations. Non-significant edges are depicted as dotted lines. (**B**) Multiplex RNA ISH/IF staining for neutrophil marker *CD66b*-IF, epithelial marker *EPCAM*-ISH, myeloid *TYROBP*-ISH, *IL1B*-ISH, and *CXCL1*-ISH and corresponding H&E images. Representative images of indicated CRC specimens (n=4 MMRd, n=4 MMRp) showing accumulations of neutrophils, *IL1B* and *CXCL1* signals at the malignant interface with the colonic lumen, often nearby dilated vessels (marked with arrows) or in necrotic regions (as indicated). Note also that neutrophils are sometimes observed directly within vessels (e.g. C103, inset). Scale bar: 50um.

**Supplemental Figure 7:**
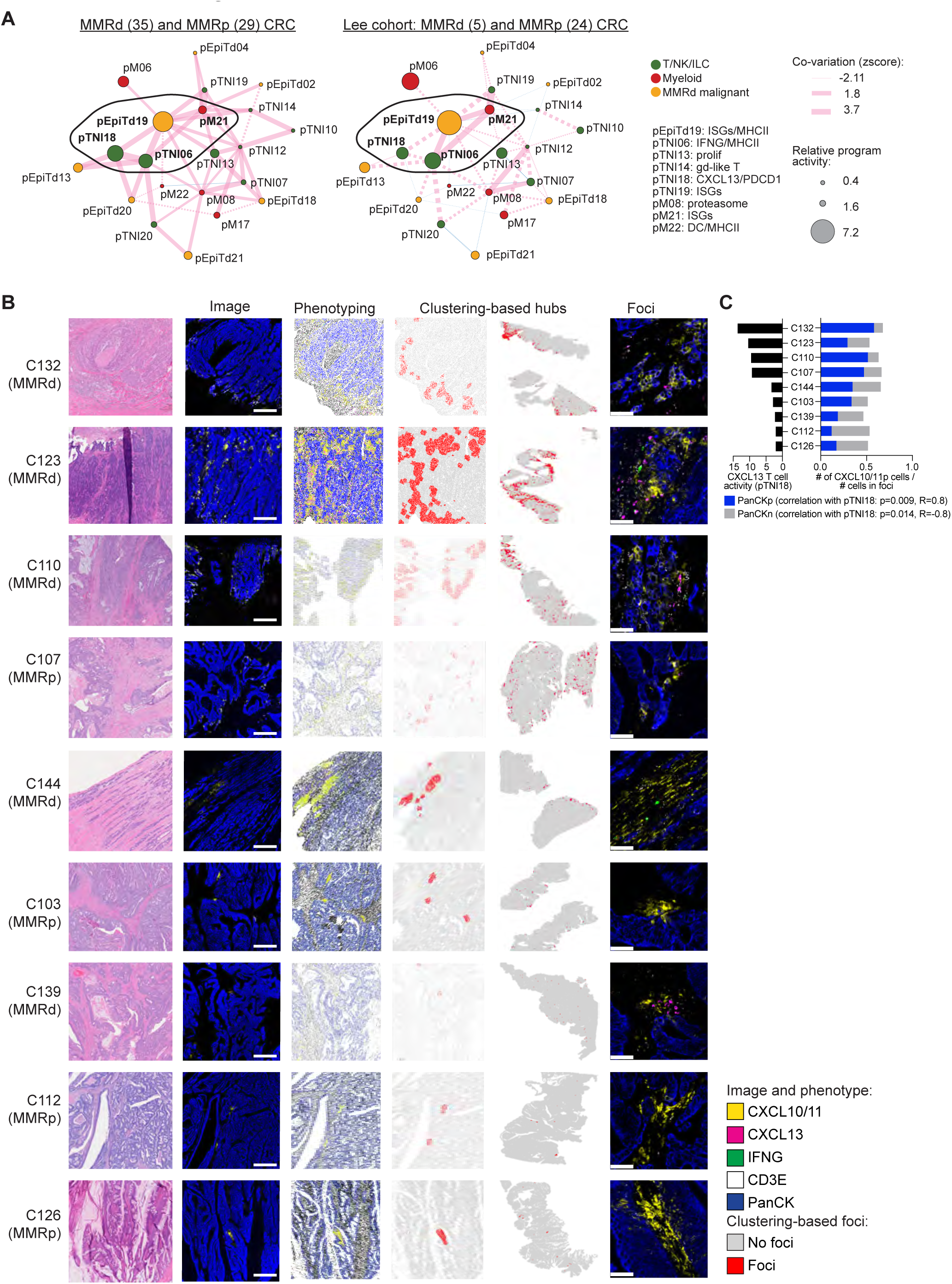
A coordinated network of *CXCL13*+ T cells, myeloid and malignant cells expressing ISGs. (**A**) ISG/*CXCL13* hub as discovered in MMRd projected onto all MMRd and MMRp specimens from our scRNA-seq cohort (left; n=35 MMRd, n=29 MMRp) or ^18^ (right; n=5 MMRd, n=24 MMRp). Node size is proportional to the log ratio of mean program activities in MMRd or MMRp vs normal. Edge thickness is proportional to co-variation scores. Pink lines depict positive, blue lines negative correlations. Non-significant edges are depicted as dotted lines. (**B**) Multiplex RNA ISH/IF staining for epithelial marker PanCK-IF, T cell marker *CD3E*-ISH, *CXCL10/CXCL11*-ISH, *CXCL13*-ISH, and *IFNG*-ISH on 9 different patient sections (MMRd n=5: C110, C123, C132, C139, C144; MMRp n=4: C103, C112, C126, C107). Cells were phenotyped using the Halo software and clustered by their neighborhoods (defined as 100 um) into cells that are part of the foci or not (red and grey, respectively). Shown from left to right for each patient specimen are an H&E section, fluorescent image, a computational rendering of the same section, the assignment to foci in the same section, the assignment of foci in the whole slide scan and magnified fluorescent images of foci. Scale bars: 500um for second column, 50um for right-most column. (**C**) For each specimen (ordered by their scRNA-seq-based *CXCL13* T cell activity) the fraction of *CXCL10/CXCL11*-positive PanCK-positive and *CXCL10/CXCL11*-positive PanCK-negative cells within foci is shown. High *CXCL13* T cell activity correlates with higher fractions of *CXCL10/CXCL11*-positive PanCK-positive cells (Spearman correlation).

## TABLES

**Supplemental Table 1.** Clinical characteristics of patient cohort, summary of 10x channels and cell subsets. Related to Figure 1.

**Supplemental Table 2.** The immune compartment in MMRd and MMRp CRC and adjacent normal colon tissue – cellular composition and transcriptional programs. Related to Figure 2.

**Supplemental Table 3.** The stromal cell compartment in MMRd and MMRp CRC and adjacent normal colon tissue – cellular composition and transcriptional programs. Related to Figure 3.

**Supplemental Table 4.** The epithelial compartment (malignant and non-malignant) in MMRd and MMRp CRC and adjacent normal colon tissue – cellular composition and transcriptional programs. Related to Figure 4.

**Supplemental Table 5.** Covariation analysis in MMRd and MMRp CRC. Related to Figures 5, 6 and 7.

**Supplemental Table 6.** In vitro stimulation of SNU-407 CRC cell line. Related to Figures 4, 6 and 7.

**Supplemental Table 7.** Imaging analysis. Related to Figures 7.

## Materials and Methods

### Lead Contact

Further information and requests for resources and reagents should be directed to and will be fulfilled by the Lead Contact, Nir Hacohen.

### Materials Availability

This study did not generate new unique reagents.

### Data And Code Availability

The single cell data set generated during this study are available at the Single Cell Portal: https://singlecell.broadinstitute.org/single_cell/study/SCP1162

All code used to analyze data is available upon request from Lead Contact.

### Human tumor specimens

Institutional Review Boards at MGH and BWH approved protocols for tissue collection used for sequencing. Informed consent was obtained from all subjects prior to collection. Age and sex of subjects can be found in **Supplemental** Table 1. Only patients with primary treatment-naive colorectal cancer were included in this study. Two patients were excluded after collection when it was discovered that they had concurrent hematologic neoplasms: myelofibrosis/AML (C108) and multiple myeloma (C117). Patient H&E slides were from the pathology department archives.

## Experimental Methods

### Establishment of in vitro fibroblast culture

A primary fibroblast culture was derived from a human CRC organoid culture established from an MMRd specimen from a 64 yo female patient. Initiation and culture of the MMRd CRC specimen was performed as described previously^73^. Fibroblasts grew out the matrigel, adhered to the bottom of the plate and were separated from the CRC organoid culture during passaging. Upon separation from CRC organoids, fibroblasts were further expanded in DMEM supplemented with 10% FBS, L-Glu, and PenStrep and frozen down in 90% FBS + 10% DMSO.

### In vitro cytokine stimulation of fibroblasts and CRC cells

Primary fibroblasts derived from CRC specimen and SNU-407 MMRd CRC cell line (male) were seeded in 96 well plate (20K cells/well fibroblasts, 50K cells/well CRC cells), rested overnight and then stimulated for 14h with 10 ng/ml IL6, TNF, IL1A, or IL1B, or left untreated. Upon stimulation, cells were lysed in TCL with 1% BME (50 ul per well). Smart-seq2 protocol was used as previously described^74^ to generate mini-bulk RNA-seq libraries (with 500 cells starting material per condition). Libraries were sequenced on Illumina NextSeq500 Sequencer. Results are representative of two independent experiments. SNU-407 cell line was fingerprinted at the Broad Genomic Platform, to make sure SNPs match the original line, and tested for mycoplasma.

### Tissue processing, CD45 enrichment, and scRNA-sequencing

Samples were cut by pathology assistants at MGH and BWH hospitals. To preserve the invasive border for clinical pathological evaluation, the pathology assistants did not sample tumor down to the invasive border. The tissue was transported in ice cold RPMI with 5% human serum prior to processing. Tissue was transferred into a petri dish on ice. Fat, necrotic, and fibrous areas were removed. Residual blood and stool were washed off the tissue with cold RPMI. Tissue allocated for dissociation was minced into small pieces (∼1 mm^3) using a scalpel prior to enzymatic dissociation. Thereafter tissue was transferred into 1.5 ml Eppendorf tubes, each containing 1 ml of enzymatic digestion mix (Miltenyi, Human Tumor Dissociation kit). 1 ml of digestion mix was used per 50 mg of tissue. The Eppendorf tubes were then transferred to a rotation shaker set to 37 °C and 550 rpm and shaken for 20 min. The digestion mix was subsequently filtered through a 70um cell strainer sitting on a 50 ml falcon tube on ice and mechanically dissociated once more with the plunger of a 1ml syringe against the screen. The filter and enzymatic mixture were washed with RPMI containing 2% human serum as needed until the suspension passed through the filter. The cell suspension was spun at 500 g for 7 min in 4 °C pre-cooled centrifuge to pellet the cells. The pellet was lysed in 4ml ammonium-chloride-potassium (ACK) buffer for 2 minutes and then stopped with RPMI containing 2% human serum. The cell suspension was then centrifuged at 500 g for 7 min at 4 °C. The resulting cell pellet was resuspended in loading buffer (PBS containing 0.04% m/v BSA) and filtered through the cell strainer snap cap (Corning 352235) into a 1.5 ml Eppendorf tube. The cell suspension was centrifuged at 500 g for 2 min at 4 °C. The pellet was resuspended in cold loading buffer (PBS containing 0.04% m/w BSA) and counted by trypan exclusion. The suspension was then diluted to 1000 cells / ul. 8000 cells were loaded into each channel of the 10x Chromium controller, following the manufacturer-supplied protocol for the 3’ kit. Additionally, a CD45-enriched sample was run for each specimen. To this end, dissociated and ACK-lysed cells were resuspended in cold PBS with 2 mM EDTA and 0.5% FCS and CD45+ cells were enriched using CD45 MicroBeads, human (Miltenyi) following the manufacturer’s instructions. Cells were resuspended in loading buffer and loaded with 8000 cells per channel as described above. 10x libraries were constructed using the 10x supplied protocol and sequenced at the Broad Institute Genomics Platform.

### RNAscope in situ hybridization with co-immunostaining

Patient cohort for RNAscope analysis was: C103 (MMRp), C107 (MMRp), C110 (MMRd), C112 (MMRp), C123 (MMRd), C126 (MMRp), C132 (MMRd), C139 (MMRd), C144 (MMRd). 5um sections were cut from formalin-fixed paraffin-embedded blocks onto SuperFrost plus slides and baked at 65 °C for 2 hours before use. Mixed RNAscope (Advanced Cell Diagnostics)/antibody antigen retrieval and staining with Opal (Akoya Biosciences) fluorophores was performed on a Leica Bond Rx instrument following the RNAscope LS Multiplex Fluorescent v2 Assay combined with Immunofluorescence protocol (322818-TN). The only two variations from the written protocol were (1) an open wash dispense after the peroxide step and (2) DAPI (Sigma D9542) was dispensed twice at the end of the protocol at a concentration of 1ug/mL. Slides were rinsed in water (Fisher 23-751628) prior to coverslipping (Fisher 12-544C) with mountant (Life Technologies P36961). Stained slides were imaged using a Vectra Polaris microscope.

### Nanostring GeoMx® Digital Spatial Profiling method to measure the expression of **∼**1500 genes in paired epithelial and non-epithelial regions

5um formalin-fixed paraffin-embedded tissue sections were baked at 65 °C for 1 hour and manually prepared using the manufacturer supplied V1.4 protocol (MAN-10087-03). Per protocol, the slides were washed thrice for 5 minutes in CitriSolv, and then twice for 5 minutes in each of 100% ethanol, 95% ethanol, and then water. Antigen was retrieved by placing slides in a staining jar containing 1x Tris EDTA (pH 9) and incubated at low pressure at 100 °C for 20 minutes. This was followed by a 5 minute wash in PBS. Thereafter, slides were placed in a staining jar with 1ug/mL proteinase K and incubated at 37 °C for 15 minutes. After proteinase digestion, slides were washed in 10% neutral buffered formalin for 10 minutes. This step was followed with two washes in a stop buffer containing tris and glycine and one wash in 1x PBS. The RNA probe mix (pre-commercial version of Cancer Transcriptome Atlas probeset) was diluted in buffer R and this hybridization solution was pipetted over the tissue, covered with a hybrislip coverslip, and incubated overnight at 37 °C. The following morning, the coverslips were removed and slides washed twice with a stringent wash containing SSC and formamide at 37 °C and then twice with SSC. The slides were then stained with fluorescently labeled morphology markers (CD45, Pan-cytokeratin, CD8, and Syto13) for 1 hour and then washed twice in SSC.

Slides were loaded on the GeoMx® microscope for imaging and barcode acquisition, following the manufacturer supplied protocol (MAN-10102-01). An overview scan at 20x was acquired. 45 circular regions of interest measuring 500um in diameter were placed on slides. ROIs were segmented into PanCK-positive and -negative areas of interest. The digital mirrored display was then employed to direct the UV laser to collect barcodes according to the specified collection masks.

Library preparation was performed according to manufacturer instructions (Nanostring DSP-Genomics Library Preparation Protocol 01/2019). Per protocol, a PCR mastermix and well-specific indices were employed to index and amplify the collected wells in a thermocycler. Thereafter, amplified barcodes were pooled and purified using AMPure XP beads and ethanol washes. A Bioanalyzer DNA high sensitivity trace was used to assess library quality. Samples were sequenced on the NextSeq2000 platform.

### Computational Analysis

#### scRNA-seq pre-processing and quality control filtering

For droplet-based scRNA-Seq, CellRanger v3.1 was used to align reads to the GRCh38 liftover (37 liftover, v28) human genome reference. The output was processed using the dropletUtils R package (version 1.7.1), to exclude any chimeric reads that had less than 80% assignment to a cell barcode, and identify and exclude empty cell droplets^75, 76^, by testing against a background generated from barcodes with 1,000 to 10 UMIs, with cutoffs determined dynamically based on channel specific characteristics. UMI and gene saturation was estimated in individual cells by sub-sampling reads without replacement in each cell barcode, in incremental fractions of 2%, with 20 repeats. A saturation function of the form 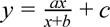 was fit based on the number of UMIs observed while sampling reads at different depths. Cell barcodes were excluded if they satisfied any one of the following criteria: (**1**) Fewer than 200 genes; (**2**) Fewer than 1,000 reads; (**3**) Fewer than 500 UMIs; (**4**) More than 50% of UMIs mapping to the mitochondrial genome; (**5**) Non-empty droplet with false discovery rate (FDR) less than 0.1; or (**6**) Over 5% of reads estimated as coming from swapped barcodes/chimeric reads (available at the Single Cell Portal, see **Data Availability**). The filtered data was clustered and cells were manually assigned to immune/stromal/epithelial groups based on expressed markers. Using outlier exclusion separately for each channel and each channel cell-type combination, cells that deviated by >2 interquartile ranges (IQR) from the median were then flagged based on the following criteria: (**1**) log_10_(total transcript UMI), (**2**) Fraction of barcode swaps, (**3**) Gene saturation estimate, (**4**) UMI saturation estimate, (**5**) Fraction of UMI supported by >1 read. Cells were further flagged if they substantially deviated from the fit based on the following relationships: (**1**) Total reads vs total UMI, (**2**) Total UMI vs log likelihood of being empty; (**3**) Total UMI vs total number of genes. A cell was excluded if it was flagged by at least two of these criteria for epithelial and immune cells, or at least three criteria for stromal cells.

#### Selection of variable genes, dimensionality reduction and clustering

After filtering and exclusion, scRNA-seq profiles were clustered across all patients using a non-negative matrix factorization (NMF) and a graph clustering-based approach. Transcriptionally over-dispersed genes were identified within each experimental batch (*i.e.*, 10x channel) by the difference of the coefficient of variation (CV) from the median CV for other genes with a similar mean expression^77^. A robust set of 1,000 to 8,000 genes was retained based on an elbow-based criterion, applied to the median of over-dispersed difference statistic based on 200 samples of 75% of cells. Next, 80% of genes and samples were sub-sampled between 50 to 200 times, and NMF was used to reduce the dimensionality of the full dataset to between 15 and 40 dimensions as the product of two non-negative matrices^24^. The loading matrices (*i.e.*, activations) of these NMF components were used to calculate *k*-nearest neighbors (*k*-NN) graph (*k*=21) based on a cosine similarity distance. This graph was clustered using stability optimizing graph clustering (http://michaelschaub.github.io/PartitionStability/,^78, 79^), to identify 7 top level cell type clusters (epithelial, stromal, mast, B, plasma, myeloid, and T cells). The same procedure was then applied iteratively to the cell profiles of each top-level cell-type separately, resulting in a total of 88 cell clusters spanning distinct types or states (**Supplemental** Table 1).

#### Cluster connectivity

To identify relationships between clusters (‘cluster connectivity’) we used Partition-based Graph Abstraction (PAGA) with connectivity model v1.2 on the NMF based *k*-NN graph above. PAGA edge thresholds were selected by using the minimum edge weight of the corresponding minimum spanning tree for each *k*-NN graph (Figure 3D-F).

#### Cluster assignment by gradient boosting and filtering of potential doublets

In order to exclude potential doublets and low confidence assignments by clustering we used a classifier for final assignment of cells to clusters. Gradient boosting (R 3.6.1, xgboost v0.90.0.2) was first applied to build a cell to cluster classifier for each of the top-level seven cluster types and subsequently to each of the 88 low-level clusters. During training, we restricted to include only the highest quality cells: (1) we excluded potential doublets, defined as cells appearing by manual examination between major high-level cell-type regions with expression features from both cell types; (2) cells with possible quality concerns that were not substantial enough for removal during QC; (3) cells with elevated potential ambient RNA contamination, retaining 314,524 cells (85%) for final classifier training.

For each of the seven top-level cell-types, a separate classifier was trained to predict each cell type separately (one-vs-all), in a 5-fold cross-validation scheme. Next, using the probability scores of the held-out test-set we identified an optimal cutoff for each class based on an ROC analysis comparing the true positive rate (TPR = true positives divided by all positive predictions) to the false positive rate (FPR = true negative divided by all negatives) and selecting the point at which the ROC curve intersects with the diagonal. Cells that were ambiguously assigned in this way to more than one cluster were removed as potential doublets.

Next, a similar classification training scheme was applied separately to cells from each top-level cell-type (epithelial, stromal, mast, B, plasma, myeloid, and T cells). We used 5-fold cross-validation and ROC analysis to select thresholds. In cases where a cell was assigned to more than one subtype, we used the assignment with the higher predictive score. Cells that could not be assigned confidently by any classifier were excluded from further analysis.

#### Classifying malignant cells by gradient boosting

Adjacent normal tissue, which was sampled distantly from the tumor (*e.g.*, ∼10cm apart), is expected to be tumor-cell free. We used gradient boosting to train a classifier predicting malignant from non-malignant epithelial cells based on the source channel type (tumor vs. adjacent normal), in a 5-fold cross validation scheme. We separately trained two classifiers, one predicting isTumor and another predicting isNormal, and used the geometric mean of the resulting probabilities as the final statistic. In subsequent analyses, we considered epithelial cells from tumor channels with a predicted score greater than 0.75 to be malignant, and cells from normal channels with a predicted score <0.25 to be normal epithelial cells. Overall, by this measure ∼95% of tumor channel epithelial cells were predicted to be malignant, and 98% of normal channel epithelial cells were predicted non-malignant cells. The classifier predictions were highly concordant with those made by inferred copy number alterations with only ∼11% of likely malignant cells showing no substantial copy number differences from normal (8% for MMRp, and 15% for MMRd), and 2% of likely normal cells showing copy number differences (data not shown). Copy number alterations were only determined for epithelial cells.

#### Identification of novel gene expression programs by NMF

To identify robust transcriptional programs, we adapted a consensus NMF procedure^22^. We used as input the weight components matrices (W matrices) from an NMF procedure that was run on 50-200 subsampled gene x cell subsets, as described above (see section on **Selection of variable genes, dimensionality reduction and clustering**). We excluded outlier components by sorting components by their cosine distance to the 20^th^ nearest neighbor and excluding components with unusually high distance by an elbow-based criterion. Next, we constructed a *k*-NN graph (*k*=30), and identified clusters of highly similar components in this graph using stability optimizing graph clustering^78^, with an exponentially varied scale parameter (0.1 to 10). The components in each cluster were median-averaged into a single component, resulting in a shortlist of “consensus NMF” components. These were used as the initialization component matrix for a second round of NMF of all cells and highly variable genes (as described in **Selection of variable genes, dimensionality reduction and clustering**). The above procedure was applied separately to each top-level cell population and to epithelial cells from normal channels. For each cell type, this resulted in eight solutions, of between 8-48 clusters corresponding to different choices of the resolution parameter. For each cell type, a single solution was selected based on examination of the mean cluster silhouette, inflection of residual error graph, and by manual examination of the top genes in the output programs.

To characterize the novel expression programs identified with this procedure, we used the top 150 genes in each of the components, ranked by the following weighting scheme: For the *i*^th^ gene and *j*^th^ component we define the scaled weight as follows: 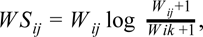 where *^W^ik* is the largest weight for gene *i* in the rest of the components, i.e., *k* ≠ *j*. This weighting scheme prioritizes for high weight (highly expressed; first term in *WS_ij_* formula) and unique genes in each component (second term in *WS_ij_* formula).

#### Identification of shared gene programs in malignant epithelial cells

To identify expression programs shared across malignant epithelial cells, from multiple individual patients, the above consensus NMF procedure was first applied separately to malignant cells from each patient (cells from tumor channels and classified as malignant as described above). For each patient a separate consensus NMF expression program set (*W* matrix) is generated, with the number of programs chosen automatically based on the residual error graph. Next, a similar consensus approach was applied to the combined list of all per-patient consensus NMF program sets (all W matrices, one per patient) as well as a set of 17 normal epithelial programs (identified as described above **Identification of novel gene expression programs by NMF),** in order to capture malignant and normal epithelial programs in a single combined NMF solution. After this consensus clustering procedure had completed, NMF clusters including one or more normal epithelial programs were excluded and the corresponding normal NMF programs were used in their place. This was done for all specimens (resulting in 43 pEpi programs), and separately for MMRd and MMRp tumors (resulting in 29 pEpiTd and 32 pEpiTp programs, respectively, **Supplemental** Table 2**-4**).

#### Calculating NMF transcriptional program activity

In order to calculate the NMF program activity matrix (*H*), we used non-negative least squares (NNLS), solving the following equation for the matrix *H*, *H* = *argmin_H_*_>0_|*X* − *WH*|*_F_*, given *X* and *W*, where *H* is the ‘program activity’ matrix, *k* is the by cell matrix; *X* is the gene by cell expression matrix, and *W* is the gene by *k* NMF expression program matrix. *W* was restricted to at most top 100 weighted genes per NMF component (selected as described above - **Identification of novel gene expression programs by NMF**). In this way we can calculate the activity values for any cell including cells not part of the original NMF procedure used to discover the program “dictionary” (e.g. pEpiTd* in MMRp cells or in data from^18^).

#### Testing for enrichment of TF targets in transcriptional programs

We tested the set of top genes from each transcriptional program for enrichment of TF target genes based on TF targets taken from http://www.regnetworkweb.org/home.jsp and estimated significance with the hypergeometric test.

#### Testing for covarying NMF expression programs

We calculate the covariation of two programs A, B as the correlation (see below) between the vectors of their program activity across the patients, where program activity is calculated by the cell type in which the program was initially defined (e.g. pTNI* programs in T/NK/ILC cells). We restricted this analysis to include only patients where at least 1,000 cells were captured and did not consider stromal cells due to their relative low fraction (<5% of all profiled cells). In order to capture relationships between expression programs that are active in only a small number of cells, we calculate for each patient, cell type, and expression program, the program activity values in this cell type at five quantiles (0.25 0.5, 0.75, 0.95, 0.99). We then calculate the Pearson correlation coefficient, *R,* for every pair of NMF programs, separately for each quantile across patients. The correlation for each quantile was Fisher transformed (i.e. arctanh(R)) and the mean of the five values was used as a test statistic and compared against a null distribution of mean Fisher transformed R values generated by permuting the patient ID assignment (and keeping cell type, and overall NMF value distribution unchanged). A p-value was calculated by counting how often the permuted R is above the true observed R (P = (# R>R’)/(# permutations), and separately how often the permuted R is below the observed R. The minimum of these (scaled by two) was taken as the outcome empirical p-value statistic and reported at a Benjamini-Hochberg FDR of 10%. We report the raw correlation at the 0.75 quantile and the adjusted R, calculated as the difference of mean true R values, and the mean of permuted R values across 10,000 permutations. We constructed a signed weighted network from the pairwise R values retaining only 288 significant edges (FDR<0.1).

Next, we discovered modules (‘hubs’) in the resulting network using a module detection algorithm for signed graphs (i.e., having both negative and positive edges,^80^). This method explores a space of solutions set by a resolution parameter in the range 0.001 to 0.2, and a random-walk parameter (tau=0.2), and outputs the optimal solution based on the Constant Potts Model of graph modularity. We applied this method iteratively, and split modules if they were larger than 3 nodes and improved the signed weighted modularity of the solution.

#### Constructing a network of expression programs similarity

A network of expression program similarities was constructed for pTNI*, pS*, pM*, and pEpi* programs by calculating for every pair of program genes a pairwise Jaccard similarity (i.e. for sets A and B J = |A intersect B|/|A union B|) of the top 50 program genes (selected as described above - **Identification of novel gene expression programs by NMF**). The resulting similarity matrix was used to construct a Gaussian kernel matrix (as in constructing a tSNE, with perplexity of 30 and a tolerance of 10^-5^). The kernel matrix was filtered to retain the top 4% of value pairs to construct the final network, and visualized using a force-directed layout algorithm.

#### Visualization of single cell profiles

We generated tSNE plots per compartment from NMF loading matrices, with a perplexity value of 30 and the Barnes-Hut approximation method^81^. A global tSNE of all cells was generated using Pegasus with the default parameters and using SVD for the preliminary embedding (v0.17.0,^82^).

#### Identification of differentially expressed genes

Differentially expressed genes (DEGs) were identified using a two-step procedure applied to the log_2_(UMI count/10^4^ + 1) values, first using a Mann-Whitney-Wilcoxon Ranksum test, and then sorting genes by Wilcoxon statistic, and testing each of the top 1,000 genes for differential expression using a generalized linear mixed model using a normal distribution, with terms for the total UMI and the total number of genes, and a fixed effect intercept term for each patient. We report the likelihood ratio Wald-test p-value comparing this model to one also including a categorical class term.

Genes were identified as differentially expressed in a particular set of cells if they met all of the following criteria: (**1**) Rank sum test with a Benjamini-Hochberg FDR < 0.1; (**2**) Minimum expression in at least 5% of cells; (**3**) Area Under a Receiver Operating Curve (AUROC) > 0.55, (**4**) 1.25 log fold change *vs.* all other cells; and **(5)** Wald-test with a Benjamini-Hochberg FDR < 0.1. We included tables for the top 100 significant genes (sorted by AUCROC), for immune, stromal and epithelial cells (**Supplemental** tables 2-4).

#### Pearson residuals calculation in contingency tables

The Pearson residual is a measure of relative enrichment for cells in a contingency table. It is calculated here as: 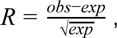 where the expected value is calculated as the product of row and column marginal probabilities by the total count.

#### Preprocessing of bulk RNA-seq data from fibroblast and cancer cell line stimulation experiment

Reads were extracted from image files using bcl2fastq2 (v2.20.00). 2×67nt paired-end reads were mapped to the human genome (GRCh37liftOver) using STAR v2.7.3a and TPM (transcripts per million) was calculated with RSEM v1.3.1. The resulting matrix was log_2_(x+1) transformed for downstream analysis.

#### Preprocessing of microarray datasets

Microarray datasets were downloaded from GEO (GSE39582: https://www.ncbi.nlm.nih.gov/geo/query/acc.cgi?acc=GSE39582,^31^; GSE13294: https://www.ncbi.nlm.nih.gov/geo/query/acc.cgi?acc=GSE13294,^32^) and pre-processed in R to match probe IDs to gene symbols according to the specified microarray chip platform “[HG-U133_Plus_2] Affymetrix Human Genome U133 Plus 2.0 Array” with chip definition table GPL570 (https://www.ncbi.nlm.nih.gov/geo/query/acc.cgi?acc=GPL570). For genes represented by multiple probes, the mean value of all probes was taken.

#### Preprocessing of bulk RNA-seq from TCGA

Standardized RNA-seq expression data for TCGA-COADREAD (CRC) samples was downloaded from GEO along with clinical annotation tables (GSE62944,^83^). We used log(TPM) values for downstream analysis.

#### Calculating gene signature scores in bulk expression datasets

We calculated gene signature scores to assess NMF program activities and fibroblast clusters in external bulk RNA-seq cohorts and ISG/MHCII scores in NanoString GeoMx data (Figure 7). We used the AddModuleScore function of the *Seurat* v3 R package^84, 85^. For each sample, this calculates the average expression of genes in the module subtracted by the average expression of a randomly selected set of control genes with similar expression across the samples. As input to the function, we used normalized expression as described above, and in each case, we used 200 random control genes.

For the NMF program scores, we used the top 150 weighted genes in each program (see **Supplemental** Table 2**-4**). Gene signatures for fibroblast clusters (Figure 3K) were:

- cS27 (CXCL14 CAF): CXCL14, AGT, NSG1, MEST, EMID1, CST1, BMP4, WNT4, INHBA
- cS28 (GREM1 CAF): COL10A1, GAS1, RSPO3, COL11A1, FAP, INHBA
- cS29 (MMP3+ CAF): MMP10, CCL20, IL1B, CSF2, STC1, INHBA
- Fibro all: C1S, LUM, DCN, RARRES2, COL1A2, C1R, COL6A2, COL3A1, MMP2, FBLN1, SERPINF1, COL6A1, COL6A3, COL1A1, CTSK, TMEM176B, MFAP4, SPON2, PDGFRA, TMEM176A, PCOLCE, CFD, VCAN, TIMP1, AEBP1, LGALS3BP, EMILIN1, LRP1, NUPR1, OLFML3, MEG3, FTL, CCDC80, NBL1, FTH1, CD63, LTBP4, IGFBP6, TIMP2, CLEC11A, CST3, ECM1, IGFBP5, MRC2, SDC2, PLTP, CXCL14, EFEMP2, RHOBTB3, RP3-412A9.11

Gene signature for MHCII/ISG was (Figure 7D, E):

- ISGscore nanostring: HLA-DMA,HLA-DMB,HLA-DPA1, HLA-DQB1, PSMB10, PSMB8, PSMB9, TAP1, TAP2, TYMP, STAT1, CXCL10, CXCL11, GBP1, GBP2, GBP4.

#### Image analysis with HALO

Raw Vectra Polaris images for each slide were unmixed with inForm software (Akoya Biosciences), using an algorithm built on a library of fluorescence spectra measured using single fluorophore labeled control slides. The unmixed multi-layer image TIFFs from single fields of view were then stitched together fused into a single multi-layer pyramidal TIFF in Halo software (Indica Labs). Tumor regions were manually annotated in Halo. The luminal margin was defined as the region 360 um radially out into the tumor from the line of outermost growth toward the lumen, and any tissue radially into the lumen was included in the luminal margin. Areas of low tissue quality such as folds, tears, bubbles, edge artifacts, and necrotic tissue were excluded. The FISH-IF v1.2.2 Halo module was used for cell segmentation and phenotyping. The resulting object dataframe was used for calculating phenotypic composition and for further neighborhood and cluster analysis (described in **Image analysis, neighborhood definition, and clustering**). With the exception of very highly expressed genes, ISH fluorescence was dot-like. The minimum unit dot area and intensity to define a copy were empirically determined by a pathologist (JHC). Copies were recorded as a semi-quantitative measure of expression in the output dataframe. Copies were also binned into categories in accordance with recommendations from Advanced Cell Diagnostics: 0, 1+ (1-3 copies/cell), 2+ (4-9 copies/cell), 3+ (10-15 copies/cell), and 4+ (>15 copies/cell). All ISH probes were called positive if they were category 1+ or above, with the exception of the secreted factors *CXCL1*, *IL1B*, and *RSPO3*, which were categorized as positive if they were 4+.

#### Image analysis, neighborhood definition, and clustering

For each full slide microscope image, the object data generated with HALO was used to extract a neighborhood for each cell. The neighborhood was defined as all cells within 100 micrometers (um) and was characterized by: 1) the total number of cells in the neighborhood; 2) the number of cells in the neighborhood from each of the following phenotypes: PanCK+, CXCL10/CXCL11+, CXCL13+, IFNG+, CD3E+, CD3E+IFNG+, CD3E+CXCL13+, PanCK+CXCL10/CXCL11+, PanCK+CXCL10/CXCL11-, AllNeg; 3) the mean and median distances to each of the cellular phenotypes, where the distance was set to 150um if no cells of a given phenotype were found in the neighborhood; 4) the sum and max of the ‘Copies’ feature for each ISH stain: CXCL10/CXCL11+, CXCL13+, IFNG+, CD3E+.

To identify ‘immune-foci’ vs ‘non-foci’ areas we used k-means clustering to cluster cells into two clusters (kmeans() functions from R *stats* package v4.0.1 with parameters: k=2, nstart=10, iter.max=10), where each cell was represented by the sum and max ‘Copies’ features of its neighborhood. To ensure that clustering results are comparable across all 9 MMRp and MMRd images, the data from all images was concatenated and clustered simultaneously. The cluster with fewer cells was labeled as the foci-cluster, which was validated by manual examination in all 9 images. We also performed *k*-means clustering after shuffling the cell ID-to-neighborhood mapping and ensured that the percent of cells assigned to cluster 2 (i.e. considered foci) for the 9 images was significantly lower (p=0.003906, Wilcoxon signed rank exact test):

| C132 | C123 | C110 | C144 | C139 | C107 | C126 | C112 | C103 |
| --- | --- | --- | --- | --- | --- | --- | --- | --- |
| 3.25 | 8.83 | 3.22 | 0.47 | 0.15 | 2.12 | 0.15 | 0.06 | 0.33 |
| 0.06 | 2.40 | 0.03 | 0.00 | 0.00 | 0.02 | 0.00 | 0.00 | 0.00 |

The total number of cells per image and numbers of cells within or outside of foci are recorded in **Supplemental** Table 7.

